# Emergent robustness of bacterial quorum sensing in fluid flow

**DOI:** 10.1101/2020.10.23.352641

**Authors:** Mohit P. Dalwadi, Philip Pearce

## Abstract

Bacteria use intercellular signaling, or quorum sensing (QS), to share information and respond collectively to aspects of their surroundings. The autoinducers that carry this information are exposed to the external environment; consequently, they are affected by factors such as removal through fluid flow, a ubiquitous feature of bacterial habitats ranging from the gut and lungs to lakes and oceans. To understand how QS genetic architectures in cells promote appropriate population-level phenotypes throughout the bacterial life cycle requires knowledge of how these architectures determine the QS response in realistic spatiotemporally varying flow conditions. Here, we develop and apply a general theory that identifies and quantifies the conditions required for QS activation in fluid flow by systematically linking cell- and population-level genetic and physical processes. We predict that, when a subset of the population meets these conditions, cell-level positive feedback promotes a robust collective response by overcoming flow-induced autoinducer concentration gradients. By accounting for a dynamic flow in our theory, we predict that positive feedback in cells acts as a low-pass filter at the population level in oscillatory flow, allowing a population to respond only to changes in flow that occur over slow enough timescales. Our theory is readily extendable, and provides a framework for assessing the functional roles of diverse QS network architectures in realistic flow conditions.

## I. INTRODUCTION

Bacteria share and respond collectively to information about their surrounding environment through the production, release and detection of small diffusible molecules called autoinducers, in a process termed quorum sensing (QS). In QS systems, the individual bacterial expression of genes relevant to the community is promoted when autoinducers accumulate to a threshold concentration, typically associated with an increasing cell density [1]. Population-level behaviors exhibited in QS-activated states include bioluminescence [2, 3], virulence factor production [4], modified mutation rates [5], biofilm and aggregate formation [6, 7], and biofilm dispersal [8]. As autoinducers diffuse between cells, they are often subject to complex and fluctuating features of their environment, such as extracellular matrix components [9, 10], interference by other bacterial species (or the host organism), and external fluid flow. Recent research has started to show how such environmental factors are closely linked to the QS response, building on foundational knowledge gained from studying well-mixed laboratory cultures [11–13]. However, improving our understanding of the functional role of QS systems requires understanding how these systems promote appropriate population-level phenotypes in realistic bacterial environments.

Fluid flow is ubiquitous in a diverse range of bacterial habitats from rivers, lakes, and medical devices to the host teeth, gut, lungs, and nasal cavity [14]. In addition to its mechanical effects on the structure of cell populations [15–19], external fluid flow has been found to have a strong influence on the transport of relevant chemicals including nutrients [8, 20], antibiotics during host treatment [21, 22], and QS autoinducers [23–26]. Recent experimental [23–27] and numerical [28–34] studies suggest that flow-induced autoinducer transport can affect population-level phenotypes by introducing chemical gradients within populations and, if the flow is strong enough, suppressing QS altogether. These results raise two important questions about QS genetic networks. Firstly, how can QS networks ensure a robust population-level response in order to avoid individual cells committing to a costly multicellular phenotype in isolation, while also avoiding premature population-level QS activation in a spatiotemporally complex environment? Secondly, how can QS networks enable populations to sense cell density in flow environments that promote high mass transfer [35–38]?

Here, we answer these questions by combining simulations and a systematic asymptotic analysis of QS in a cell layer subject to an external flow; we focus on the effect of positive feedback in autoinducer production, a common feature of QS genetic circuits [39]. First, we establish the conditions required for the emergence of population-level QS activation in steady flow. Our results illustrate how the required conditions for activation depend on the ratio of the timescale of the external flow to the timescale of diffusion through the cell layer. If the required conditions are met in a region of the cell layer, positive feedback causes autoinducers to flood the population, inducing population-wide QS activation. Interestingly, by accounting for a dynamic flow in our model we find that an ability to avoid premature QS activation is built into systems with positive feedback. We predict that positive feedback acts as a low-pass filter to oscillations in the shear rate; if such oscillations occur over a time period shorter than a critical time that we calculate, the QS system is not activated, even if the required conditions for activation are met during the oscillations. Furthermore, we find that by combining multiple QS signals, a population can infer both cell density and external flow conditions. Overall, our findings suggest that positive feedback allows QS systems to act as spatiotemporally non-local sensors of fluid flow.

## II. RESULTS

### Population-level theory for QS in flow

To understand how genetic circuits in individual cells affect population-level bacterial signaling, we investigated an archetypal quorum sensing (QS) circuit in Gram-negative bacteria called a LuxIR system (Fig. 1a; Methods). In this system, autoinducers (AI) in cells bind to a cognate LuxR protein and the bound AI-LuxR dimer promotes the transcription of downstream genes. The system exhibits a positive feedback loop through the presence of an AI synthase, LuxI, whose expression is promoted by the bound dimer [12, 39]. Thus, to summarize, as the concentration of AI in a cell increases, there is an increase in the number of bound dimers. Consequently, this promotes the production of LuxI, which further increases the production of AI.

**FIG. 1:**
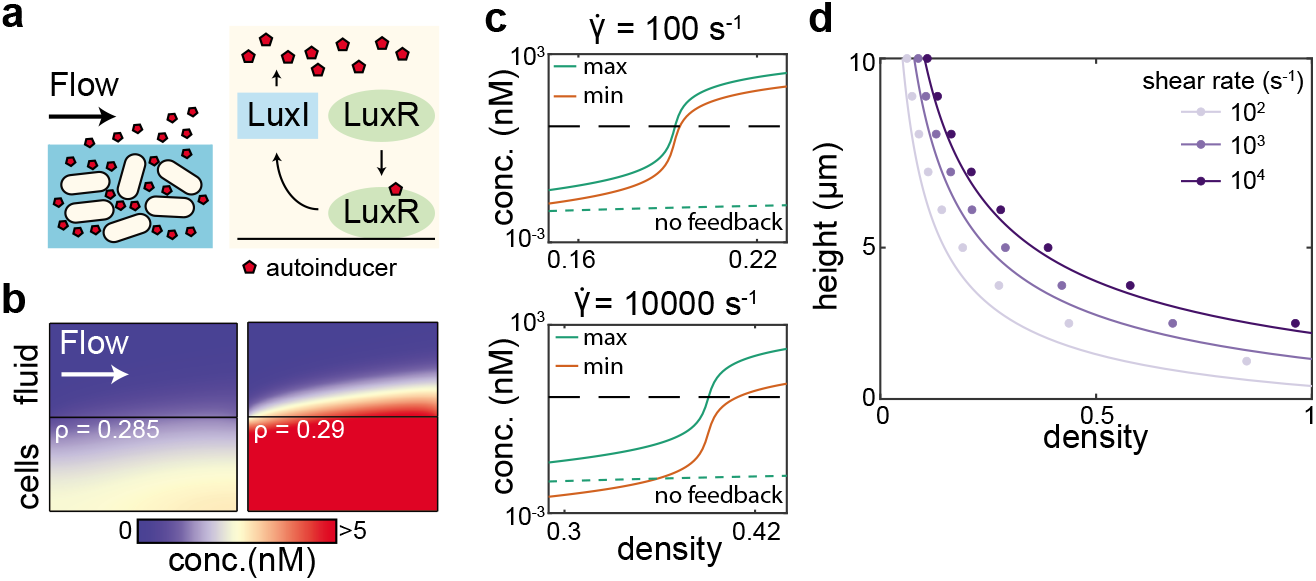
We identify the required conditions for the onset of quorum sensing (QS) activation in a LuxIR-type system in steady flow. a) We model a population of cells, and an external fluid flow above the population (left). Inside cells, we model a LuxIR-type genetic circuit with positive feedback [12, 39] (right). b) In steady simulations for an imposed uniform shear flow, if the density is below a critical value, the system remains inactivated (left). If the density rises just above this critical value, positive feedback causes robust population-level QS activation (right). c) The maximum (green line) and minimum (red line) autoinducer concentrations inside a population rise drastically around the critical density in steady simulations. Results are shown for shear rates of 100 s^−1^ (top) and 10000 s^−1^(bottom); the population size was taken to be *H* = 5 *μ*m. Dashed black lines correspond to the activation threshold, and dashed green lines show the maximum concentration in the cell population when no feedback is present (λ = 0); we note that setting the binding parameters *k*_+_ = 0 and *k*_-_ = 0 does not have a distinguishable effect in that case. d) The simulations show that the critical density is larger for a larger shear rate, and smaller for a larger cell population (dots). Lines show the predicted critical density from Eqs. (6)–(8). All kinetic parameters in these simulations are given the values listed in Table S1.

We modeled the concentration of AI (A), LuxR (*R*), LuxI (*I*), and bound AI-LuxR dimers (*C*) in a population of cells through a locally-averaged set of governing equations [40, 41]. The important cell-scale information is captured through the local volume density of cells *ρ* (Methods). We consider the scenario where cells are embedded in extracellular matrix (matrix generation precedes QS activation in species including *Vibrio cholerae* [8] and *Pseudomonas aeruginosa* [25]) over which a fluid flows. As such, the fluid flow imparts a shear stress (which may vary in space and time) to the upper boundary of the cell layer. We neglect growth-induced flows inside the cell layer because their timescales are typically much slower than those of diffusion and external flow [18, 42].

Thus, our problem consists of two coupled domains. To obtain non-dimensional equations in each domain, we scale lengths with the height of the cell layer *H*, fluid velocities with 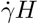, where 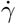 is a typical shear rate, and times with the diffusion timescale *H*^2^/*D_c_*, where *D_c_* is the AI diffusion coefficient in the cell population (see Methods). In the cell population region, a diffusion-reaction equation holds for the AI concentration

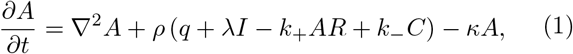

where ∇^2^*A* represents AI diffusion, *q* is the base production rate of AI, λ is the synthesis rate of AI by LuxI, *k*_+_ is the binding rate of AI and LuxR proteins, *k*_-_ is the corresponding unbinding rate, and *κ* is the decay rate of AI. Reaction equations hold for the concentrations of the proteins and dimers inside cells

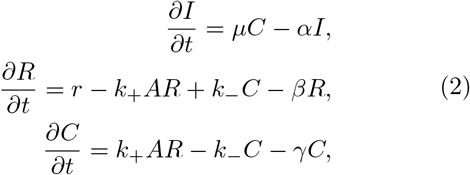

where *μ* is the activation rate of LuxI by AI-LuxR complexes, *r* is the base production rate of LuxR, and *α, β* and *γ* are decay rates. We combine the kinetic parameters into two parameter groups,

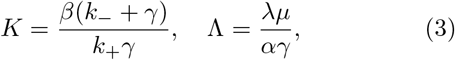

where *K* represents an effective equilibrium constant for AI-LuxR complex formation, and Λ represents the strength of positive feedback in the system. In the external flow region, with flow field ***u***, an advection-diffusion equation holds for the AI concentration

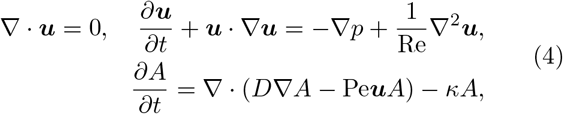

where Re is the Reynolds number of the flow, and *D* is the ratio of AI diffusivity in the flow to AI diffusivity in the cell layer. The key control parameter that we use to investigate how external flow affects QS in cell populations is the Peclet number

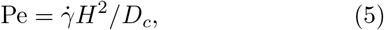

which quantifies the relative effects of advection in the external flow to diffusion in the cell layer.

### QS activation in steady flow

First, we performed numerical simulations of the governing equations in the finite-element computational software COMSOL Multiphysics for an imposed spatially uniform shear flow, which we assumed to be free of AI far upstream (see Fig. S1 for a full description of the numerical procedure). This incorporates a very wide range of laminar flows in simple geometries owing to the large Schmidt number (Sc = *ν/D_e_* > 1000, where *ν* is the kinematic viscosity and *D_e_* is the diffusion coefficient in the flow) of AI in water, so that the flow profile can be linearized in the mass-transfer boundary layer. We performed steady simulations to understand the conditions in which it is possible for a population to enter a QS-activated state for typical kinetic and physical parameter values (summarized in Table S1). The results show that a strong flow can entirely suppress QS activation by removing AI from the population boundary (Fig. 1b). However, above a critical cell density *ρ_c_*, the cell population is able to exhibit QS activation through the positive feedback present in the system. At steady state in this regime, the domain becomes flooded with AI, which increase in concentration by several orders of magnitude throughout the population (Fig. 1b); the large increase does not occur in systems without positive feedback (Fig. 1c). This change occurs over very small changes in cell density (Fig. 1c). In larger cell populations, populations with restricted or reduced AI diffusion, and weaker external flows, the critical density is smaller owing to the reduced mass transfer of AI out of the population (Fig. 1d, Fig. S5).

To understand the general principles that guide how the various kinetic, physical and geometric parameters determine *ρ*_c_, we analyzed the system of equations for a thin cell layer, where diffusion through the cell layer in the direction of flow is much less important than diffusion in the direction normal to the surface of the cell layer [43]. In this systematically reduced model, the entire effect of the external flow region on the AI concentration *A* within the cell population is reduced to an effective Robin boundary condition on the surface of the cell layer. To derive this condition, we constructed a similarity solution for *A* in the mass-transfer boundary layer [44] in the external fluid (see Supporting Information). This yielded an effective Péclet number, Pe_eff_, which quantifies the local ratio of advective to diffusive transport at the position *x* (where the x-axis is directed with the flow and *x* = 0 corresponds to the upstream edge of the population). Our analysis predicts the effective Péclet number to be

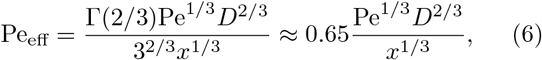

where *D* is the ratio of the diffusion coefficient in the external flow to the diffusion coefficient in the cell layer (see Eq. 16). Our Pe_eff_ prediction agrees well with simulation results (Fig. 2a) outside a small diffusive boundary layer of thickness O(Pe^−1/2^) at the downstream end of the cell population (see Fig. S3).

**FIG. 2:**
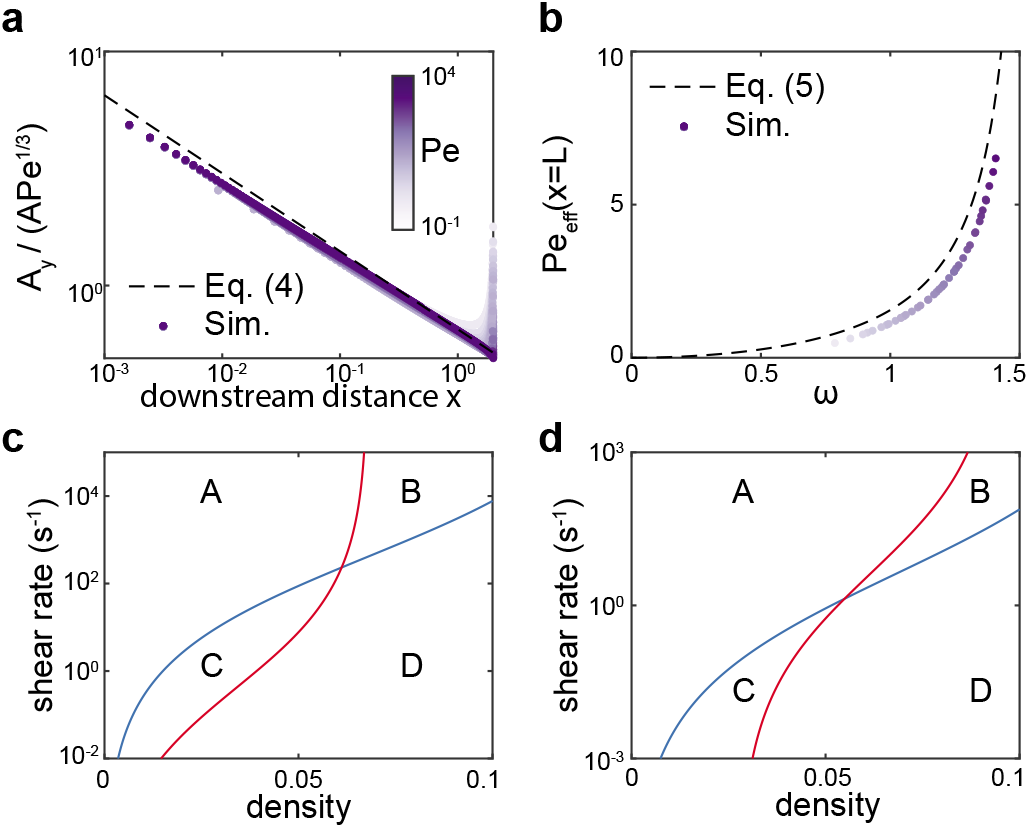
We identify and analyze an algebraic relationship linking flow, biomass and kinetics at the onset of quorum sensing (QS) activation. a) For the simulations in Fig. 1d, when the calculated effective Péclet number at the cell population boundary, *A_y_/A*, is scaled by Pe^1/3^, the simulation data collapse onto the curve 0.65/*x*^1/3^ (dashed line), as predicted in Eq. (6). Here A is the autoinducer (AI) concentration, *x* is the co-ordinate parallel to the flow, and *y* is the co-ordinate perpendicular to the flow. b) When plotted against the appropriate non-dimensional variables identified in Eqs. (6)–(7), the simulations from Fig. 1d collapse onto the curve defined by Eq. (7) (see also Fig. S4). c,d) Illustrative examples of how cells can measure density and flow by measuring activation (Eq. 7) of two different AI, splitting parameter space into four regions A, B, C and D. c) Cell population of height 10 *μ*m; one AI has kinetic parameters from Table S1 (blue line), and the second AI also has a factor of 10 reduction in diffusivity in the cell layer and factor of 5 reduction in LuxR production rate (red line) compared to the other AI. d) Cell population of height 100 *μ*m; one AI has kinetic parameters from Table S1, but with factor of 10 increase in LuxI and LuxR decay rates (blue line), and the second AI also has a factor of 500 increase in AI decay rate and factor of 2 increase in LuxR production rate (red line) compared to the other AI.

The steady thin-film governing equations admit a solution for *A* that satisfies an ordinary differential equation, which we analyze through the method of matched asymptotic expansions (see Supporting Information). Based on the expected orders of magnitude of the parameter values (see Table S1), we exploit the physiologically relevant limits *K* ≫ 1 and Λ ≫ 1, corresponding to a relatively large equilibrium constant for AI-LuxR complex formation, and strong positive feedback, respectively (see Eq. 3). Our analysis demonstrates that, for a fixed Pe_eff_, the system exhibits an imperfect transcritical bifurcation at a critical density, which marks an orders of magnitude increase in *A* owing to a drastic increase in positive feedback. We identify this point as the critical density *ρ_c_* above which the population exhibits QS activation, which reveals the algebraic relationship

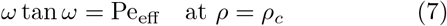

where

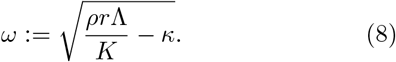

Here *r* and *κ* represent LuxR production and AI decay, respectively (see Eqs. 1–2). In Eq. (7), the effect of flow is captured in Pe_eff_ and the effect of the kinetic parameters at the population level is captured in the single non-dimensional parameter group *ω*.

To compare the predictions from our systematically reduced model to the simulation results, we note that the onset of QS activation will occur at the lowest effective Péclet number at the boundary of the cell population. Because Pe_eff_ decreases in *x* (see Eq. 6), we ignore the small downstream boundary layer for simplicity and assume that Pe_eff_ is minimized at the downstream end of a cell population of length *L*. Therefore, we insert *x* = *L* into Eq. (6) to predict Pe_eff_ for a general population of cells in a uniform shear flow. Plotting our simulation results against the non-dimensional parameter groups Pe_eff_ and *ω* (defined in Eqs. 6 and 8, respectively) demonstrates a remarkable collapse onto the predicted curve Eq. (7) for a wide range of typical kinetic, geometric and physical parameters (Fig. 2b, Fig. S4), despite the assumptions in our thin-film reduction. We note that we slightly underestimate Pe_eff_ (and therefore *ρ_c_*) due to the thin diffusive region at the downstream end of the population; an improvement would require a full spatial asymptotic analysis of the problem. This collapse onto Eq. (7) can be ‘unwrapped’ to calculate the critical conditions for activation for a QS network with a given set of kinetic parameters.

For example, we found that, by combining two different AI signals, a bacterial population can respond separately to the cell density and the external shear rate, by measuring the activation state of both signals. The key tunable parameters for sensing this difference are the diffusion coefficients of each AI within the cell layer, the decay rates of each AI, and the kinetics of each QS network through the strength of the positive feedback (or, more specifically, the ratio *r*Λ/*K* in Eq. 7). We found that two of these parameters must be different between the two AI signals to separate parameter space into four regions that correspond to low and high values of both cell density and shear rate (Fig. 2c,d). In thin cell layers, where the overall decay of AI within the cell layer is very small for typical parameters (see Table S1), populations can combine two signals with different diffusion coefficients and different strengths of positive feedback (Fig. 2c). In larger cell layers, populations can also incorporate information from two AI with different decay rates (Fig. 2d). We note that in very large populations, our model may need to be modified to account for nutrient limitations (see Fig. S10). This ability to measure cell density and shear rate separately is possible because the three tunable parameters affect QS activation conditions in distinct ways: AI diffusivity within the cell layer has a larger effect at larger shear rates, AI decay has a consistent effect across shear rates, and the effect of positive feedback is strongly dependent on the cell density.

### QS activation in complex geometries

In complex geometries there will be regions of low shear on the cell layer surface, on which the local effective Péclet number Eq. (6) will be reduced. Eq. (7) predicts that the global *ρ_c_* will be lowered in such cell populations, in agreement with a recent experimental study in which QS activation was found to be promoted in crevices or pores [26]. To further understand the local and global effects of complex geometries on QS, we performed simulations of the 3D governing equations in channels which mimic typical host environments such as intestinal crypts and tooth cavities. The channels contain crevices that extend in the horizontal direction transverse to a pressure-induced flow over a cell population that coats the channel floor (Fig. 3a; Fig. S1). We found that *ρ_c_* is reduced in channels with crevices, by an amount that depends on the crevice depth (Fig. 3b) and the number of crevices (Fig. S6), in agreement with experimental findings [26]. Furthermore, we found that once QS is activated in the crevices, diffusion of autoinducers activates further regions of the cell population outside the crevices, particularly downstream (Fig. 3c). This activation region can extend for lengths far beyond the size of the crevices themselves, even in conditions for which QS activation would be precluded completely in a simple channel (Fig. 3b,c). This demonstrates that local geometric complexities can have highly non-local effects on QS activation through positive feedback.

**FIG. 3:**
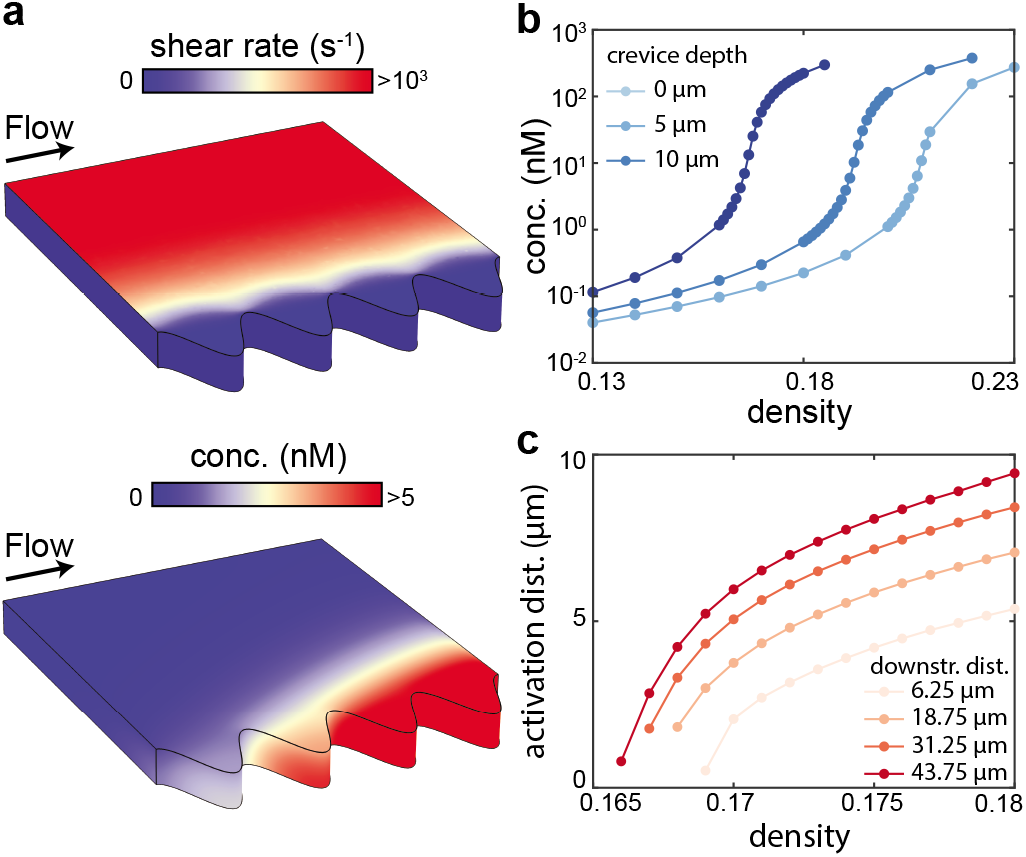
In complex geometries, positive feedback causes robust QS activation based on the region with the lowest effective Péclet number. a) We simulated steady flow in a channel with crevices in its sidewall; a cell population coats the channel floor. Top: The shear rate applied by the flow to the top of the cell population is lower in the crevices (no shear is applied to the sides of the cell population). Bottom: The autoinducer concentration in the cell population is higher in the crevices and downstream. Kinetic parameters correspond to those in Table S1, with *ρ* = 0.168 and a crevice depth of 10 *μ*m, which is twice the height of the population (see Fig. S1 for further details of the simulation geometry and boundary conditions). b) Steady simulations show that QS is activated at lower densities in populations in channels with deeper crevices. The maximum autoinducer concentration in the population, which occurs in the most downstream crevice (see panel a), is plotted. c) Steady simulations show that for a larger density, a larger region of the population is activated. The density is plotted against the QS-activated distance along the line transverse to the center of each of the four crevices; the downstream distance of each crevice from the upstream edge of the population is labeled. The QS-activated region is defined as the region for which *A* > 5 nM, which we take to be the activation threshold. All kinetic parameters were given the values listed in Table S1.

### QS activation in unsteady flow

To understand the transient process of QS activation in unsteady flows, which are common in bacterial habitats such as the lungs and medical devices, we performed simulations of the dynamic governing equations for spatially uniform flows with a sinusoidally oscillating shear rate. Each oscillating flow is characterized by the time *t*_act_ during an oscillation period for which the system is in the ‘QS activation region’ of parameter space,

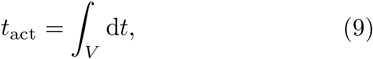

where *V* is the set of times such that Pe_eff_(*t*) < Pe_eff_(*t_c_*) over one oscillation, and *t_c_* is the time at which the effective Péclet number Pe_eff_ falls below its critical value identified in Eq. (7) (see inset of Fig. 4a). For the range of shear rates spanned by the oscillations, the steady solution for AI concentration varies over orders of magnitude as the system passes through the critical Pe_eff_ (Fig. 4a). However, in the dynamic simulations, for a fixed mean and amplitude of oscillation, this range is only achieved for long enough oscillation periods (i.e. larger *t*_act_). Surprisingly, for shorter oscillation periods (i.e. smaller *t*_act_), the AI concentration remains below the QS-activation threshold; we note that throughout these simulations, *t*_act_ remains well above the diffusive timescale. We found that there is a critical oscillation period (i.e. a critical *t*_act_), and that as the period increases over this critical value, the mean (and maximum) concentration of AI increases over several orders of magnitude (Fig. 4b, Movie S1). We did not observe such a critical oscillation period for a system without feedback (λ = 0), suggesting that this effect is caused by positive feedback (Fig. S8).

**FIG. 4:**
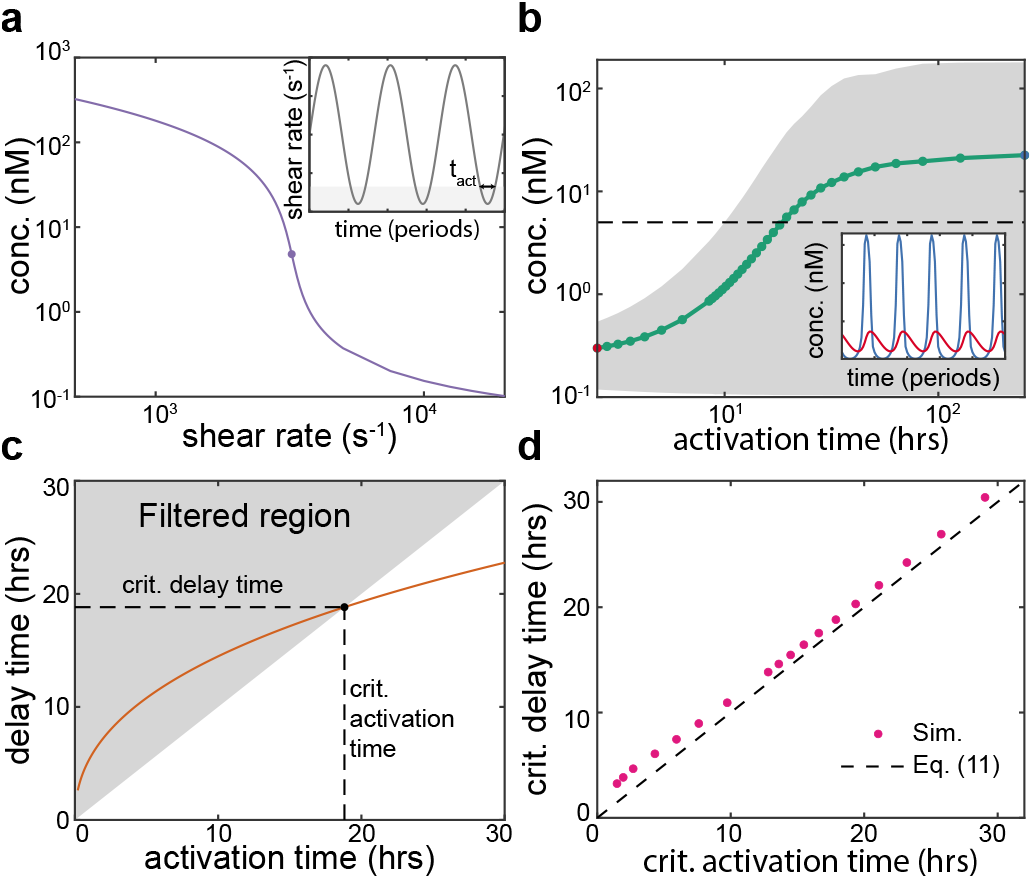
We identify the required conditions for QS activation in an oscillating flow. a) In steady simulations with fixed density, the maximum concentration of autoinducers rises drastically when the shear rate is below its critical value. In the following simulations we oscillate the shear rate across this critical value with a fixed mean and amplitude; the oscillations are defined by the activation time *t*_act_ for which the system is in the QS activated region of parameter space (inset). b) Simulations were performed with oscillations in the shear rate with mean 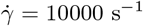 and amplitude 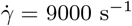. For oscillation time periods slower than a critical time, corresponding to a critical *t*_act_, the mean (green line) and range (grey area) of the maximum autoinducer concentration throughout each oscillation rise drastically (see also Fig. S7 and Movie S1). Oscillations in the autoinducer concentration are filtered out for *t* < *t*_act_, but not for *t* > *t*_act_ (inset). c) Over a range of oscillation periods, for each oscillation we calculate the activation time using Eq. (9) and the delay time using Eq. (10). Thus, we predict a critical oscillation period by finding the oscillation at which the activation time is equal to the delay time. If the activation time is below the delay time, the oscillation is in the filtered region of parameter space, and the QS system is predicted to remain inactivated. d) To confirm the validity of our prediction of the critical oscillation period, we performed 1D simulations of the thin-film equations, where the Péeclet number is directly controllable. We defined the critical oscillation as the one at which the maximum autoinducer concentration first rose above the threshold value of 5 nM; then, we calculated the critical activation time using Eq. (9). We found that, when plotted against the predicted critical delay time (Eq. 10), this critical activation time collapses onto Eq. (11). All kinetic parameters in these simulations were given the parameter values in Table S1.

To explore how this observation depends on the LuxIR system kinetics, and to determine whether it is a general property of the system, we performed a dynamic analysis of the thin-film equations as Pe_eff_ passes below its critical value at *t* = *t_c_* and the system enters the ‘QS activation region’ of parameter space. We found that there is a critical slowing down due to the imperfect transcritical bifurcation which marks the onset of QS activation (see Supporting Information); these dynamics are reminiscent of the effects of ‘ghosts’ of saddle-node bifurcations [45–47]. This slower timescale introduces a ‘delay time’, *t*_delay_, which determines the time *t* = *t_c_* + *t*_delay_ at which dynamic QS activation occurs if the effective Péeclet number remains below its critical value, i.e. if Pe_eff_(*t*) < Pe_eff_(*t_c_*) for *t* > *t_c_*. The delay time depends on the system kinetics through two non-dimensional parameter groups *ν*_1_ and *ν*_2_, (defined in Eq. 17; see Methods), and on the imposed external flow through the time-derivative 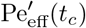 of the effective Péclet number as it passes below its critical value:

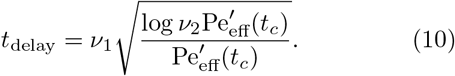

Thus, the delay time depends on whether the shear rate changes slowly or quickly through the critical value that marks the onset of QS activation.

Interestingly, this result suggests that in a dynamic flow, even if the shear rate falls below its critical value into the QS activation region, the onset of QS activation would not be triggered if the shear rate increases back above its critical value after a time shorter than the delay time of the system. Therefore, for an oscillating flow that enters the QS activation region of parameter space for an activation time *t*_act_ during each oscillation, we identify the oscillation period as the critical period if the activation and delay times are equal (Fig. 4c):

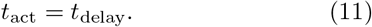

Eqs. (10)–(11) predict that for a flow oscillating over a long enough time period, such that *t*_act_ > *t*_delay_, the QS system passes through cycles of dynamic activation and deactivation. Conversely, for a flow oscillating over a short enough time period, such that *t*_act_ < *t*_delay_, the system is predicted to remain in the QS-inactivated state. We confirmed the validity of our prediction of the critical oscillation period by performing simulations of the governing thin-film equations, in which the Péclet number and its derivatives are directly controllable. The results show that for a wide range of oscillating Péclet numbers, the calculated properties for the onset of dynamic QS activation collapse onto the curve defined by Eqs. (10)–(11) (Fig. 4d).

In experiments, and in our simulations of the governing equations with an imposed flow, it is not possible to control the effective Péeclet number and its derivatives directly. However, we can make an order of magnitude estimate of the required conditions for the onset of dynamic QS activation for a sinusoidally oscillating flow with a time period *T_p_* as follows. At onset, 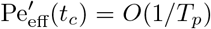 and *t*_act_ = *O*(*T_p_*). Combining Eq. (10) and (11) and neglecting the effect of the logarithmic term in Eq. (10) (which we expect to have a lesser effect than the algebraic terms) yields an estimate of 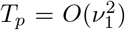 at the onset of dynamic QS activation. For the typical kinetic parameters used in our simulations (see Table S1), this suggests a critical oscillation period of approximately 10 hours, which is in agreement with our simulation results for a wide range of oscillating flows (Fig. 4b,d; Fig. S7).

## III. DISCUSSION

This study demonstrates how positive feedback in the LuxIR system, an archetypal bacterial quorum sensing (QS) genetic circuit, promotes a robust population-level response in spatiotemporally varying flow conditions. Because QS systems measure the concentration of passively transported autoinducers (AI), even simple fluid flows generate concentration gradients which can cause phenotypic gradients within a population. Our results show that positive feedback in QS genetic architectures allows bacteria to overcome flow-induced AI gradients at the population level for a wide range of conditions that represent flows encountered in bacterial habitats such as lakes, rivers, and hosts (Figs. 1, 3). The key physical determinant of the onset of QS activation in a population is the minimum value of the effective Péeclet number Pe_eff_ (see Eq. 6), which quantifies the local advective to diffusive transport at the surface where the population meets the external fluid. Furthermore, a systematic reduction of the governing equations via a thin-film model reveals that for a given Pe_eff_, the critical population density (or size) for the onset of QS activation is determined by a single dimensionless kinetic parameter group *ω* (see Eq. 8). A compact relationship between these two parameter groups, Eq. (7), links the physical, geometric and kinetic parameters at the onset of QS activation (Fig. 2). Through their transparent dependence on these system parameters, Eqs. (6)–(8) explain how QS activation is promoted in bacteria with QS architectures with stronger positive feedback or with smaller AI-LuxR dissociation constants [48]; in conditions of restricted AI diffusion inside the population, which can be caused by interactions with the extracellular matrix [9] (Fig. S5); in larger or denser populations; and in populations subject to weaker external flow, in agreement with recent experimental results [25, 26].

The dependence of the critical density on the flow conditions raises the question of whether bacterial QS systems in fluid flow respond to increasing cell density, decreasing mass transfer, or a combination of these factors [37]. By calculating the QS activation conditions for AI with different sets of physical and kinetic parameters using Eqs. (6)–(8), we suggest that a bacterial population can integrate information from multiple signals to measure cell density and shear rate separately (Fig. 2c,d). Our results are in qualitative agreement with previous work which considered a well-mixed population subject to spatially uniform AI removal by mass transfer and AI decay [37]. However, our results also suggest that in smaller populations subject to external flow, the overall decay of AI may be too small to separate parameter space into distinct regions; in such populations, physical differences between AI diffusivities (which could be caused by different interactions with surrounding matrix proteins [9]) may provide more information. These results suggest that by combining multiple AI signals with different physical and kinetic properties, bacterial populations in complex environments can add fidelity to measurements of their surrounding conditions, and promote the appropriate phenotypic response to these conditions.

An individual bacterium committing to a QS-activated phenotype can incur significant individual costs, such as the generation or abandonment of important extracellular material. It is therefore often beneficial for such a commitment to be shared by the rest of the bacterial population [49]. As we have shown, positive feedback promotes a robust, population-scale QS response if appropriate conditions are met in only a local region of a cell population. While this feature of positive feedback is very useful for population-wide commitment, it has the potential downside of triggering premature QS activation in noisy or intermittent flow conditions. However, our analysis of the LuxIR system in dynamic flow conditions suggests that the positive feedback mechanism actually reduces the potential for such a premature response because the feedback itself acts as a low-pass filter at the population level. That is, in a flow oscillating with period smaller than a critical value that we calculate (but still slower than the diffusion timescale), a population responds to the mean effective Péeclet number, rather than exhibiting quasi-steady oscillations in QS activation and inactivation (Fig. 4). In such conditions, a growing population’s QS system would be expected to activate eventually through its increasing density (growth is also associated with an activation delay time but for typical parameters it is of the same order of magnitude as the doubling time; see Fig. S9). The critical time period depends on the nature of the flow oscillation and the system parameters through Eq. (10), which manifests due to a bottleneck induced by an imperfect transcritical bifurcation in the system; such critical slowing down is a universal feature of dynamical systems near critical points [45–47]. Overall, this result suggests that population-level low-pass filtering via positive feedback complements other previously identified sources of noise-filtering, or time-averaging [50], in QS systems such as slow AI-LuxR unbinding [51] and diffusional dissipation [52, 53].

Our findings allow us to interpret typical QS network features in a manner that accounts for their expected response to spatiotemporal variations in flow. Our analysis suggests that bacterial species with positive feedback in their QS network, such as *P. aeruginosa*, use QS as a spatiotemporally non-local sensor of flow conditions and cell density. In these systems, positive feedback causes AI to flood the population if the required conditions are met in a local region, inducing QS activation in a large proportion of the population. Built into this mechanism is an ability to avoid premature population-wide activation in unsteady flow through the delay time that we have identified; the required conditions must persist for long enough for activation to occur. Our predictions of the effects of flow on AI concentration in species in which feedback does not link back to AI production, such as *V. cholerae* and *V. fischeri* [39, 54], are shown in Fig. 1c and Fig. S8. In these systems, AI concentration is much less sensitive to cell density and shear rate, and if the required conditions for QS activation are met in a local region of the population, this does not cause other regions of the population to be activated. Furthermore, in unsteady flows, such systems will exhibit repeated variations in AI concentration, and these variations will occur over diffusive timescales, which are usually faster than timescales of flow variation. Therefore, we expect these species to use QS as a spatiotemporally local sensor of flow conditions and cell density.

To conclude, we have identified the required conditions for the emergence of robust, population-level QS activation in a spatiotemporally varying fluid flow, for systems that exhibit positive feedback in their QS network architecture. We have demonstrated that positive feedback allows cells to avoid an isolated or premature commitment to costly multicellular phenotypes. Furthermore, we have found that populations can integrate multiple signals to sense cell density and flow conditions separately. Our theory demonstrates how QS genetic architectures play a key role in determining the population-level functional response of bacterial intercellular signaling systems in complex environments.

## IV. METHODS

### A. Governing equations

Inside the cell population, the governing equations are

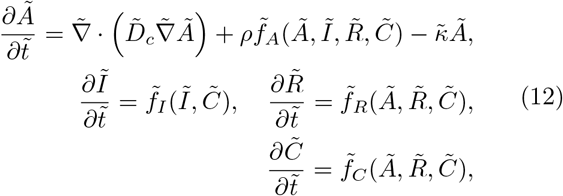

where *ρ* is the volume fraction of cells, *Ã* is the concentration of autoinducers, 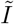 is the concentration of LuxI, 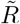 is the concentration of LuxR, and 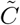 is the concentration of autoinducer-LuxR dimers. The reaction terms in the system (Fig. 1a) are

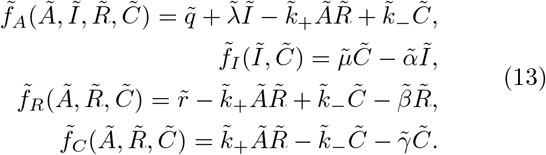

The meanings and typical orders of magnitude of each dimensional parameter are listed in Table S1. Here, for simplicity we have assumed that mRNA concentrations are quasi-steady, and have linearized the activation of LuxI by the autoinducer-LuxR dimers [55] (see Fig. S2 for a discussion of how saturation in promoter occupancy of the transcription factor affects QS activation in flow). We assume that all cells in the population have access to nutrients and are physiologically active, so that base autoinducer production 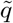 is uniform throughout the population. This assumption is based on the observation that hemispherical biofilms with radius around 10 *μ*m remain uniformly growth-active even in shear rates a hundredfold smaller than the smallest shear rates used here [17] (where a smaller shear rate implies reduced access to nutrients). This assumption may need to be relaxed in nutrient-limited conditions (see Fig. S10). Note also that we do not consider the basal expression of LuxI, which is expected to be small [56], because its only effect is to slightly change the concentration of autoinducers before activation. Outside the cell population, the governing equations are

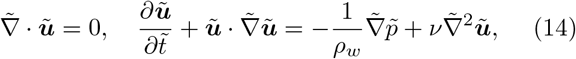

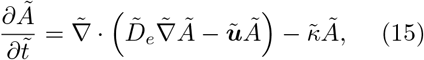

where 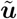 is the velocity field of the fluid, and *ρ_w_* and *ν* are the fluid density and kinematic viscosity, respectively, which we take to be those of water. We assume that the flow is free of autoinducers far upstream, and apply continuity of autoinducer concentration and concentration flux at the interface between the cell population and the flow. Full technical details of the boundary conditions, the non-dimensionalisation of the problem, and the procedures for the asymptotic and numerical solution of the governing equations are given in the Supporting Information. All code required to generate the simulation results is available on Github at https://github.com/philip-pearce/quorum-flow (requires Comsol 5.5 license, Matlab license, and Comsol LiveLink with Matlab License).

### B. Non-dimensional parameters

The non-dimensional parameters, defined in terms of the dimensional parameters in Eqs. (12)–(13), are

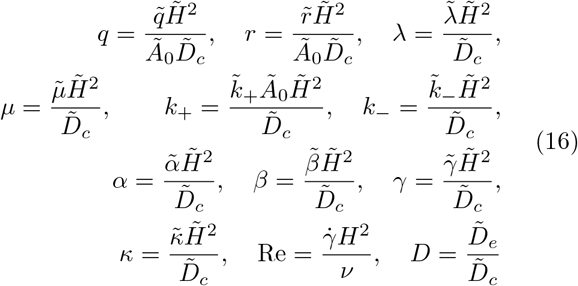

where 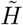 is the height of the cell population and *Ã*_0_ is the threshold concentration of autoinducers for QS activation. These parameters can be combined into the parameter groups

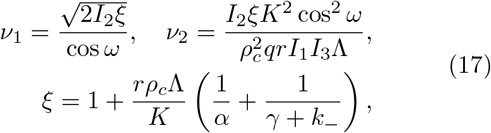

where

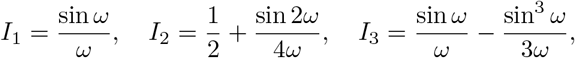

and *ω* is defined in Eq. (8).

## V. ACKNOWLEDGMENTS

We thank Johan Paulsson, Noah Olsman and Alexandre Persat for helpful discussions.

## References

[1] Papenfort, K. & Bassler, B. L. Quorum sensing signal-response systems in Gram-negative bacteria. Nature Reviews Microbiology 14, 576–588 (2016).

[2] Engebrecht, J., Nealson, K. & Silverman, M. Bacterial bioluminescence: Isolation and genetic analysis of functions from Vibrio fischeri. Cell 32, 773–781 (1983).

[3] Bassler, B. L., Wright, M. & Silverman, M. R. Multiple signalling systems controlling expression of luminescence in Vibrio harveyi: sequence and function of genes encoding a second sensory pathway. Molecular Microbiology 13, 273–286 (1994).

[4] de Kievit, T. R. & Iglewski, B. H. Bacterial Quorum Sensing in Pathogenic Relationships. Infection and Immunity 68, 4839–4849 (2000).

[5] Krašovec, R. et al. Mutation rate plasticity in rifampicin resistance depends on Escherichia coli cell–cell interactions. Nature Communications 5, 3742 (2014).

[6] Miller, M. B. & Bassler, B. L. Quorum Sensing in Bacteria. Annual Review of Microbiology 55, 165–199 (2001).

[7] Jemielita, M., Wingreen, N. S. & Bassler, B. L. Quorum sensing controls Vibrio cholerae multicellular aggregate formation. eLife 7, e42057 (2018).

[8] Singh, P. K. et al. Vibrio cholerae Combines Individual and Collective Sensing to Trigger Biofilm Dispersal. Current Biology 27, 3359–3366.e7 (2017).

[9] Charlton, T. S. et al. A novel and sensitive method for the quantification of N-3-oxoacyl homoserine lactones using gas chromatography-mass spectrometry: Application to a model bacterial biofilm. Environmental Microbiology 2, 530–541 (2000).

[10] Tarafder, A. K. et al. Phage liquid crystalline droplets form occlusive sheaths that encapsulate and protect infectious rod-shaped bacteria. Proceedings of the National Academy of Sciences 117, 4724–4731 (2020).

[11] Pérez-Velézquez, J., Gölgeli, M. & García-Contreras, R. Mathematical Modelling of Bacterial Quorum Sensing: A Review. Bulletin of Mathematical Biology 78, 1585–1639 (2016).

[12] Whiteley, M., Diggle, S. P. & Greenberg, E. P. Progress in and promise of bacterial quorum sensing research. Nature 551, 313–320 (2017).

[13] Mukherjee, S. & Bassler, B. L. Bacterial quorum sensing in complex and dynamically changing environments. Nature Reviews Microbiology 17, 371–382 (2019).

[14] Wheeler, J. D., Secchi, E., Rusconi, R. & Stocker, R. Not Just Going with the Flow: The Effects of Fluid Flow on Bacteria and Plankton. Annual Review of Cell and Developmental Biology 35, 213–237 (2019).

[15] Nadell, C. D., Ricaurte, D., Yan, J., Drescher, K. & Bassler, B. L. Flow environment and matrix structure interact to determine spatial competition in Pseudomonas aeruginosa biofilms. eLife 6, e21855 (2017).

[16] Yan, J. et al. Bacterial Biofilm Material Properties Enable Removal and Transfer by Capillary Peeling. Advanced Materials 30, 1804153 (2018).

[17] Hartmann, R. et al. Emergence of three-dimensional order and structure in growing biofilms. Nature Physics 15, 251–256 (2019).

[18] Pearce, P. et al. Flow-Induced Symmetry Breaking in Growing Bacterial Biofilms. Physical Review Letters 123, 258101 (2019).

[19] Rossy, T., Nadell, C. D. & Persat, A. Cellular advective-diffusion drives the emergence of bacterial surface colonization patterns and heterogeneity. Nature Communications 10, 2471 (2019).

[20] Stewart, P. S. Mini-review: Convection around biofilms. Biofouling 28, 187–198 (2012).

[21] Allen, R. & Waclaw, B. Antibiotic resistance: a physicist’s view. Physical Biology 13, 045001 (2016).

[22] Gralka, M., Fusco, D., Martis, S. & Hallatschek, O. Convection shapes the trade-off between antibiotic efficacy and the selection for resistance in spatial gradients. Physical Biology 14, 045011 (2017).

[23] Kirisits, M. J. et al. Influence of the hydrodynamic environment on quorum sensing in Pseudomonas aeruginosa biofilms. Journal of Bacteriology 189, 8357–8360 (2007).

[24] Meyer, A. et al. Dynamics of AHL mediated quorum sensing under flow and non-flow conditions. Physical Biology 9 (2012).

[25] Emge, P. et al. Resilience of bacterial quorum sensing against fluid flow. Scientific Reports 6, 33115 (2016).

[26] Kim, M. K., Ingremeau, F., Zhao, A., Bassler, B. L. & Stone, H. A. Local and global consequences of flow on bacterial quorum sensing. Nature Microbiology 1, 15005 (2016).

[27] Siryaporn, A., Kim, M. K., Shen, Y., Stone, H. A. & Gitai, Z. Colonization, Competition, and Dispersal of Pathogens in Fluid Flow Networks. Current Biology 25, 1201–1207 (2015).

[28] Janakiraman, V., Englert, D., Jayaraman, A. & Baskaran, H. Modeling growth and quorum sensing in biofilms grown in microfluidic chambers. Annals of Biomedical Engineering 37, 1206–1216 (2009).

[29] Vaughan, B. L., Smith, B. G. & Chopp, D. L. The Influence of Fluid Flow on Modeling Quorum Sensing in Bacterial Biofilms. Bulletin of Mathematical Biology 72, 1143–1165 (2010).

[30] Frederick, M. R., Kuttler, C., Müller, J., Eberl, H. J. & Hense, B. A. A mathematical model of quorum sensing in patchy biofilm communities with slow background flow. Canadian Applied Mathematics Quarterly 18, 267–298 (2011).

[31] Frederick, M. R., Kuttler, C., Hense, B. A. & Eberl, H. J. A mathematical model of quorum sensing regulated EPS production in biofilm communities. Theoretical Biology and Medical Modelling 8, 8 (2011).

[32] Uecke, H., Muller, J. & Hense, B. A. Individual-Based Model for Quorum Sensing with Background Flow. Bulletin of Mathematical Biology 76, 1727–1746 (2014).

[33] Zhao, J. & Wang, Q. Three-Dimensional Numerical Simulations of Biofilm Dynamics with Quorum Sensing in a Flow Cell. Bulletin of Mathematical Biology 79, 884–919 (2017).

[34] Jung, H. & Meile, C. D. Numerical investigation of microbial quorum sensing under various flow conditions. PeerJ 8, e9942 (2020).

[35] Hense, B. A. et al. Does efficiency sensing unify diffusion and quorum sensing? Nature Reviews Microbiology 5, 230–239 (2007).

[36] West, S. A., Winzer, K., Gardner, A. & Diggle, S. P. Quorum sensing and the confusion about diffusion. Trends in Microbiology 20, 586–594 (2012).

[37] Cornforth, D. M. et al. Combinatorial quorum sensing allows bacteria to resolve their social and physical environment. Proceedings of the National Academy of Sciences of the United States of America 111, 4280–4284 (2014).

[38] Fan, G. & Bressloff, P. C. Population Model of Quorum Sensing with Multiple Parallel Pathways. Bulletin of Mathematical Biology 79, 2599–2626 (2017).

[39] Ng, W.-L. & Bassler, B. L. Bacterial Quorum-Sensing Network Architectures. Annual Review of Genetics 43, 197–222 (2009).

[40] Dalwadi, M. P., Wang, Y., King, J. R. & Minton, N. P. Upscaling Diffusion Through First-Order Volumetric Sinks: A Homogenization of Bacterial Nutrient Uptake. SIAM Journal on Applied Mathematics 78, 1300–1329 (2018).

[41] Dalwadi, M. P. & King, J. R. A Systematic Upscaling of Nonlinear Chemical Uptake Within a Biofilm. SIAM Journal on Applied Mathematics 80, 1723–1750 (2020).

[42] Qin, B. et al. Cell position fates and collective fountain flow in bacterial biofilms revealed by light-sheet microscopy. Science 369, 71–77 (2020).

[43] Ward, J. P. & King, J. R. Thin-film modelling of biofilm growth and quorum sensing. Journal of Engineering Mathematics 73, 71–92 (2012).

[44] Lévêque, A. Les lois de la transmission de chaleur par convection. Annales des Mines 13, 201–299, 305–362, 381–415 (1928).

[45] Strogatz, S. H. & Westervelt, R. M. Predicted power laws for delayed switching of charge-density waves. Physical Review B 40, 10501–10508 (1989).

[46] Strogatz, S. H. Nonlinear Dynamics and Chaos with Applications to Physics, Biology, Chemistry, and Engineering (Westview Press, 2015).

[47] Gomez, M., Moulton, D. E. & Vella, D. Critical slowing down in purely elastic ‘snap-through’ instabilities. Nature Physics 13, 142–145 (2017).

[48] Sappington, K. J., Dandekar, A. A., Oinuma, K. I. & Greenberg, E. P. Reversible signal binding by the Pseudomonas aeruginosa quorum-sensing signal receptor LasR. mBio 2, e00011–11 (2011).

[49] Nadell, C. D., Xavier, J. B., Levin, S. A. & Foster, K. R. The Evolution of Quorum Sensing in Bacterial Biofilms. PLoS Biology 6, e14 (2008).

[50] Paulsson, J. Summing up the noise in gene networks. Nature 427, 415–418 (2004).

[51] Muller, J., Kuttler, C. & Hense, B. A. Sensitivity of the quorum sensing system is achieved by low pass filtering. BioSystems 92, 76–81 (2008).

[52] Tanouchi, Y., Tu, D., Kim, J. & You, L. Noise reduction by diffusional dissipation in a minimal quorum sensing motif. PLoS Computational Biology 4, 4–11 (2008).

[53] Gupta, S. et al. Investigating the dynamics of microbial consortia in spatially structured environments. Nature Communications 11, 2418 (2020).

[54] Hunter, G. A. M., Vasquez, F. G. & Keener, J. P. A mathematical model and quantitative comparison of the small RNA circuit in the Vibrio harveyi and Vibrio cholerae quorum sensing systems. Physical Biology 10, 046007 (2013).

[55] Chen, T., He, H. L. & Church, G. M. Modeling gene expression with differential equations. In Pac. Symp. Biocomput., 29–40 (1999).

[56] Goryachev, A., Toh, D. & Lee, T. Systems analysis of a quorum sensing network: Design constraints imposed by the functional requirements, network topology and kinetic constants. Biosystems 83, 178–187 (2006).

