## Supporting Information for "Emergent robustness of bacterial quorum sensing in fluid flow"

### Supporting Information: Emergent robustness of bacterial quorum sensing in fluid flow

Mohit P. Dalwadi[1] and Philip Pearce[2]
[1]*Mathematical Institute, University of Oxford, Oxford, OX2 6GG, UK*
[2]*Department of Systems Biology, Harvard Medical School, Boston, MA 02115, USA*

#### I. GOVERNING EQUATIONS

##### A. Governing equations and boundary conditions

The dimensional governing equations inside the cell layer are

$$\frac{\partial \tilde{A}}{\partial \tilde{t}} = \tilde{\nabla}\cdot\left(\tilde{D}_c\tilde{\nabla}\tilde{A}\right) + \rho \tilde{f}_A(\tilde{A},\tilde{I},\tilde{R},\tilde{C}) - \tilde{\kappa}\tilde{A}, \quad \frac{\partial \tilde{I}}{\partial \tilde{t}} = \tilde{f}_I(\tilde{I},\tilde{C}), \quad \frac{\partial \tilde{R}}{\partial \tilde{t}} = \tilde{f}_R(\tilde{A},\tilde{R},\tilde{C}), \quad \frac{\partial \tilde{C}}{\partial \tilde{t}} = \tilde{f}_C(\tilde{A},\tilde{R},\tilde{C}), \tag{S1}$$

where $\tilde{A}$, $\tilde{I}$, $\tilde{R}$, and $\tilde{C}$ are the concentrations of autoinducer, LuxI, LuxR, and autoinducer-LuxR complex, respectively, within the cell layer. Here, $\tilde{D}_c$ is the diffusivity of autoinducer within the cell layer, $\rho$ is the volume fraction of cells-to-total-volume, $\tilde{\kappa}$ is the decay rate of autoinducer, and the reaction terms $\tilde{f}_i$ (for $i \in \{A, I, R, C\}$) inside each bacterium are defined as the following:

$$\tilde{f}_A(\tilde{A},\tilde{I},\tilde{R},\tilde{C}) = \tilde{q} + \tilde{\lambda}\tilde{I} - \tilde{k}_+\tilde{A}\tilde{R} + \tilde{k}_-\tilde{C}, \tag{S2a}$$

$$\tilde{f}_I(\tilde{I},\tilde{C}) = \tilde{\mu}\tilde{C} - \tilde{\alpha}\tilde{I}, \tag{S2b}$$

$$\tilde{f}_R(\tilde{A},\tilde{R},\tilde{C}) = \tilde{r} - \tilde{k}_+\tilde{A}\tilde{R} + \tilde{k}_-\tilde{C} - \tilde{\beta}\tilde{R}, \tag{S2c}$$

$$\tilde{f}_C(\tilde{A},\tilde{R},\tilde{C}) = \tilde{k}_+\tilde{A}\tilde{R} - \tilde{k}_-\tilde{C} - \tilde{\gamma}\tilde{C}, \tag{S2d}$$

where we have linearized the term corresponding to the activation of LuxI by the autoinducer-LuxR complex (see Fig. S2).

The dimensional governing equations in the media outside the cell layer are

$$\frac{\partial \tilde{\mathcal{A}}}{\partial \tilde{t}} = \tilde{\nabla}\cdot\left(\tilde{D}_e\tilde{\nabla}\tilde{\mathcal{A}} - \tilde{\boldsymbol{u}}\tilde{\mathcal{A}}\right) - \tilde{\kappa}\tilde{\mathcal{A}}, \tag{S3}$$

where $\tilde{\mathcal{A}}$ is the concentration of autoinducer outside the cell layer, $\tilde{D}_e$ is the diffusivity of autoinducer outside the cell layer, and $\tilde{\boldsymbol{u}}$ is the fluid velocity outside the cell layer. Although we assume the autoinducer diffusivity inside and outside the cell layer is the same in the main text, we keep the two diffusivities distinct throughout the Supporting Information for generality.

At the interface between the cell layer and the and exterior medium, we have continuity of autoinducer concentration and concentration flux. Mathematically, this corresponds to

$$\tilde{A} = \tilde{\mathcal{A}}, \quad \boldsymbol{n}\cdot\tilde{D}_c\tilde{\nabla}\tilde{A} = \boldsymbol{n}\cdot\left(\tilde{D}_e\tilde{\nabla}\tilde{\mathcal{A}} - \tilde{\boldsymbol{u}}\tilde{\mathcal{A}}\right) = \boldsymbol{n}\cdot\tilde{D}_e\tilde{\nabla}\tilde{\mathcal{A}}, \tag{S4}$$

where the last equality follows from no flux of fluid into the biofilm ($\tilde{\boldsymbol{u}}\cdot\boldsymbol{n} = 0$ on the interface).

At the boundary between the cell layer and the substrate upon which it sits, we have no flux of autoinducer, corresponding to

$$\boldsymbol{n}\cdot\tilde{D}_c\tilde{\nabla}\tilde{A} = 0. \tag{S5}$$

Typical orders of magnitude of the dimensional parameters in this problem are estimated in Table S1.

##### B. Non-dimensionalisation

We scale

$$(\tilde{A},\tilde{I},\tilde{R},\tilde{C},\tilde{\mathcal{A}}) = \tilde{A}_0(A,I,R,C,\mathcal{A}), \quad \tilde{\boldsymbol{u}} = \tilde{H}\tilde{\gamma}\boldsymbol{u}, \quad \tilde{\boldsymbol{x}} = \tilde{H}\boldsymbol{x}, \quad \tilde{t} = (\tilde{H}^2/\tilde{D}_c)t, \tag{S6}$$

where $\tilde{H}$ is the typical height of the cell layer and $\dot{\gamma}$ is the typical shear rate of the fluid, so that we rescale time with the scale of vertical diffusion.

Inside the cell layer, (S1) becomes

$$\frac{\partial A}{\partial t} = \nabla^2 A + \rho\left(q + \lambda I - k_+ AR + k_- C\right) - \kappa A, \tag{S7a}$$

$$\frac{\partial I}{\partial t} = \mu C - \alpha I, \tag{S7b}$$

$$\frac{\partial R}{\partial t} = r - k_+ AR + k_- C - \beta R, \tag{S7c}$$

$$\frac{\partial C}{\partial t} = k_+ AR - k_- C - \gamma C, \tag{S7d}$$

and outside the cell layer (S3) becomes

$$\frac{\partial \mathcal{A}}{\partial t} = \nabla \cdot \left(D \nabla \mathcal{A} - \mathrm{Pe}\boldsymbol{u}\mathcal{A}\right) - \kappa \mathcal{A}, \tag{S8}$$

where we have introduced the dimensionless parameters

$$(q, r, \lambda, \mu, k_+, k_-, \alpha, \beta, \gamma, \kappa) = \frac{\tilde{H}^2}{\tilde{D}_c}(\tilde{q}/\tilde{A}_0, \tilde{r}/\tilde{A}_0, \tilde{\lambda}, \tilde{\mu}, \tilde{k}_+\tilde{A}_0, \tilde{k}_-, \tilde{\alpha}, \tilde{\beta}, \tilde{\gamma}, \tilde{\kappa}), \quad D = \tilde{D}_e/\tilde{D}_c, \quad \mathrm{Pe} = \dot{\gamma}\tilde{H}^2/\tilde{D}_c. \tag{S9}$$

On the interface between the interior and exterior, the interfacial conditions (S4) become

$$A = \mathcal{A}, \quad \boldsymbol{n} \cdot \nabla A = \boldsymbol{n} \cdot D \nabla \mathcal{A}. \tag{S10}$$

and at the interface between cell layer and substrate, (S5) becomes

$$\boldsymbol{n} \cdot \nabla A = 0. \tag{S11}$$

#### II. REDUCTION TO ONE-DIMENSIONAL PROBLEM INSIDE THE CELL POPULATION

For a thin cell layer with a small height variation, the flow near near the cell layer can be well approximated by a shear flow within the mass transfer boundary layer, assuming that the flow is laminar and the Schmidt number is large (for autoinducers in water $\mathrm{Sc} = \nu/D > 1000$, where $\nu$ is the kinematic viscosity). Choosing axes with the positive $x$-axis in the same direction as the flow, and the positive $z$-axis pointing towards the exterior media from the substrate, the biofilm interface is located at $z = 1$ at leading-order. We choose the $y$-axis to be perpendicular to both such that $(x, y, z)$ forms a right-handed Cartesian coordinate system. The fluid velocity is then

$$\boldsymbol{u} = (z - 1)\, \boldsymbol{e}_x, \tag{S12}$$

at leading-order, noting that Pe will define the strength of the flow through its dependence on the shear rate in (S9).

As the height variation in the biofilm is small, the diffusive spatial derivatives in $x$ and $y$ will be small. We work with the leading-order versions of (S7a) and (S8), which become

$$\frac{\partial A}{\partial t} = \frac{\partial^2 A}{\partial z^2} + \rho\left(q + \lambda I - k_+ AR + k_- C\right) - \kappa A \quad \text{for } 0 < z < 1, \tag{S13}$$

$$\frac{\partial \mathcal{A}}{\partial t} = D\frac{\partial^2 \mathcal{A}}{\partial z^2} - \mathrm{Pe}(z-1)\frac{\partial \mathcal{A}}{\partial x} - \kappa \mathcal{A} \quad \text{for } z > 1. \tag{S14}$$

The interfacial conditions (S10) and the biofilm-to-substrate condition (S5) become

$$A = \mathcal{A}, \quad \frac{\partial A}{\partial z} = D\frac{\partial \mathcal{A}}{\partial z} \quad \text{on } z = 1, \tag{S15a}$$

$$\frac{\partial A}{\partial z} = 0 \quad \text{on } z = 0. \tag{S15b}$$

We are interested in deriving a quasi-steady-state reduction of the governing equation outside the cell layer (S14) to an effective boundary condition at the interface of the cell layer. From steady-state numerical simulations, we find that $\mathcal{A}/x^{1/3} \approx a$ and $\partial\mathcal{A}/\partial z \approx b$ along the interface at $z = 1$, where $a$ and $b$ are approximately constant for a given set of parameter values. We also note that the timescale of autoinducer decay is much longer than the diffusive timescale (i.e. $\kappa \ll 1$). As such, we look for a similarity solution of the form

$$\mathcal{A} = x^{1/3} g(\eta), \quad \text{where } \eta = \frac{(z-1)\mathrm{Pe}^{1/3}}{(Dx)^{1/3}}, \tag{S16}$$

to balance advection and diffusion in (S14). Substituting (S16) into the steady version of (S14) and neglecting $\kappa$ results in the following ODE for $g$

$$g'' + \frac{\eta^2}{3} g' - \frac{\eta}{3} g = 0, \qquad g(0) = a, \quad g'(0) = b\left(\frac{D}{\mathrm{Pe}}\right)^{1/3}, \quad g(\infty) = 0, \tag{S17}$$

where $a$ and $b$ are the same unknown constants mentioned above. Our goal is to derive a relationship between $a$ and $b$. This will provide a relationship between $\mathcal{A}$ and $\partial\mathcal{A}/\partial z$ on the interface $z = 1$, and hence (S15a) will provide an effective boundary condition for $A$ within the cell layer, closing the interior problem.

The solution to (S17) is

$$g = \frac{a\eta}{3^{5/3}} \Gamma\left(-\frac{1}{3}, \frac{\eta^3}{9}\right), \tag{S18}$$

where $\Gamma$ is the incomplete Gamma function. Therefore, we may deduce that

$$b\left(\frac{D}{\mathrm{Pe}}\right)^{1/3} = g'(0) = \frac{\Gamma(-1/3)}{3^{5/3}} a = -\frac{\Gamma(2/3)}{3^{2/3}} a, \tag{S19}$$

and hence, using (S15a), that

$$\frac{\partial A}{\partial z} + \mathrm{Pe}_{\mathrm{eff}} A = 0 \quad \text{on } z = 1, \tag{S20a}$$

where the effective Péclet number $\mathrm{Pe}_{\mathrm{eff}}$ is defined as

$$\mathrm{Pe}_{\mathrm{eff}} := \frac{\Gamma(2/3)}{3^{2/3}} \frac{\mathrm{Pe}^{1/3} D^{2/3}}{x^{1/3}} \approx 0.65 \frac{\mathrm{Pe}^{1/3} D^{2/3}}{x^{1/3}}. \tag{S20b}$$

The effective boundary condition (S20) represents a consistent, closed boundary condition for $A$ at the interface.

#### III. REQUIRED CONDITIONS FOR QUORUM SENSING ACTIVATION IN STEADY FLOW

The coupled PDE-ODE system, systematically reduced to 1D, is given by the governing equations (S7b)–(S7d), (S13) inside the cell layer, and the boundary conditions (S15b) at the substrate interface and (S20) at the media interface.

At steady state, the three ODEs (S7b)–(S7d) result in the algebraic relationships

$$I = \frac{\mu r}{\alpha\gamma} \frac{A}{K + A}, \quad R = \frac{r}{\beta} \frac{K}{K + A}, \quad C = \frac{r}{\gamma} \frac{A}{K + A}, \tag{S21}$$

and the PDE (S13) yields the following steady-state ODE for $A$:

$$0 = \frac{\mathrm{d}^2 A}{\mathrm{d}z^2} + \rho\left(q + \frac{r(\Lambda - 1)A}{K + A}\right) - \kappa A, \tag{S22}$$

where

$$K = \frac{\beta(k_- + \gamma)}{k_+ \gamma}, \quad \Lambda = \frac{\lambda\mu}{\alpha\gamma}. \tag{S23}$$

We assume that $K \gg 1$ and $\Lambda \gg 1$, which in the context of the genetic network corresponds to a strong effect of positive feedback on the system. Under these conditions, there is a critical curve in $(\rho, \mathrm{Pe}_{\mathrm{eff}})$-parameter space across which positive feedback increases drastically; we define this curve as the conditions required for the onset of quorum sensing (QS) activation. To calculate this curve, we solve the system asymptotically in the region where positive feedback is small, and determine where the implicit asymptotic assumptions break down. This breakdown corresponds to the drastic increase of positive feedback. When positive feedback is small, $A \ll K$, and the system (S21)–(S22) is well-approximated (in the sense of matched asymptotic expansions) by

$$I = \frac{\mu r}{\alpha\gamma K}A, \quad R = \frac{r}{\beta}, \quad C = \frac{r}{\gamma K}A, \quad 0 = \frac{\mathrm{d}^2 A}{\mathrm{d}z^2} + \rho q + \omega^2 A, \quad \text{where } \omega^2 = \frac{\rho r \Lambda}{K} - \kappa \in (-\infty, \infty). \tag{S24}$$

The ODE in (S24) is solved by

$$A = \frac{\rho q}{\omega^2}\left(\frac{\mathrm{Pe}_{\mathrm{eff}}\cos\omega z}{\mathrm{Pe}_{\mathrm{eff}}\cos\omega - \omega\sin\omega} - 1\right), \tag{S25}$$

and therefore

$$\max_{z\in(0,1)} A = A|_{z=0} = \frac{\rho q}{\omega^2}\left(\frac{\mathrm{Pe}_{\mathrm{eff}}\,(1-\cos\omega) + \omega\sin\omega}{\mathrm{Pe}_{\mathrm{eff}}\cos\omega - \omega\sin\omega}\right). \tag{S26}$$

When the denominator of the fraction in the brackets vanishes, the analytic equation (S26) blows up. We note that the equation (S26) does not blow up when $\omega \to 0$, since the numerator of the fraction in the brackets also vanishes at this point. This apparent blow up is a contradiction of the small-$A$ asymptotic assumption used to derive the equation and it marks a significant increase in positive feedback in the system. As such, we may deduce the following critical curve for $\rho$ and $\mathrm{Pe}_{\mathrm{eff}}$, which we define as the onset of QS activation:

$$\mathrm{Pe}_{\mathrm{eff}} = \omega\tan\omega = \sqrt{\frac{\rho r\Lambda - \kappa K}{K}}\tan\sqrt{\frac{\rho r\Lambda - \kappa K}{K}}. \tag{S27}$$

It is also helpful to note that if $A \gg K$, once we are well into the large positive feedback regime, then

$$\max_{z\in(0,1)} A = A|_{z=0} \sim \frac{\rho r\Lambda}{\kappa}\frac{\sqrt{\kappa}\sinh\sqrt{\kappa} + \mathrm{Pe}_{\mathrm{eff}}\,(\cosh\sqrt{\kappa} - 1)}{\mathrm{Pe}_{\mathrm{eff}}\cosh\sqrt{\kappa} + \sqrt{\kappa}\sinh\sqrt{\kappa}}, \tag{S28}$$

again in the sense of matched asymptotic expansions. This marks an increase of a factor of $O(\Lambda^2/K)$ in $\max A$ across the onset of QS activation.

#### IV. REQUIRED CONDITIONS FOR QUORUM SENSING ACTIVATION IN UNSTEADY FLOW

We now consider the dynamic onset of quorum sensing (QS) activation for the 1D coupled PDE-ODE system, in the scenario where the flow is able to vary in time. The system consists of the governing equations (S7b)–(S7d), (S13) inside the cell layer, and the boundary conditions (S15b) at the substrate interface and (S20) at the media interface.

We consider the scenario where the flow varies over a timescale $t = T/\varepsilon$, with $\varepsilon \ll 1$ and $T = O(1)$. Physically, this corresponds to a timescale of flow *variation* that is slower than the timescale of vertical diffusion. In our system, the effect of flow manifests through the effective Péclet number $\mathrm{Pe}_{\mathrm{eff}}$, and QS activation is marked by decreasing through a critical effective Péclet number defined by (S27).

##### A. Away from the critical point

Over the new slower timescale $T = \varepsilon t$, the dynamical 1D system becomes

$$\varepsilon\frac{\partial A}{\partial T} = \frac{\partial^2 A}{\partial z^2} + \rho\,(q + \lambda I - k_+ AR + k_- C) - \kappa A, \tag{S29a}$$

$$\varepsilon\frac{\partial I}{\partial T} = \mu C - \alpha I, \tag{S29b}$$

$$\varepsilon\frac{\partial R}{\partial T} = r - k_+ AR + k_- C - \beta R, \tag{S29c}$$

$$\varepsilon\frac{\partial C}{\partial T} = k_+ AR - k_- C - \gamma C, \tag{S29d}$$

with boundary conditions

$$\frac{\partial A}{\partial z} = 0 \quad \text{on } z = 0, \qquad \frac{\partial A}{\partial z} + \mathrm{Pe}_{\mathrm{eff}}(T)A = 0 \quad \text{on } z = 1. \tag{S30}$$

From the form of (S29)–(S30), we see that as $\mathrm{Pe}_{\mathrm{eff}}$ changes in $T$, the dynamic system is quasi-steady. Hence, before the drastic increase in positive feedback, the asymptotic solution to (S29)–(S30) in the limit $\varepsilon \to 0$ is given by (S24)–(S25).

##### B. Near from the critical point

However, this solution breaks down near the QS activation curve, which occurs at $T = T_c$, where $\mathrm{Pe}_{\mathrm{eff}}(T_c) = \mathrm{Pe}_c$, defined by (S27). Near this breakdown, the dynamics become more important to the system. To explore these dynamics, we introduce a new timescale

$$T = T_c + \frac{\widehat{T}}{K^{1/2}}, \tag{S31}$$

where the scaling choice of $1/K^{1/2} \ll 1$ can be justified *a posteriori.* This means that the effective Péclet number can be written asymptotically as

$$\mathrm{Pe}_{\mathrm{eff}} \sim \mathrm{Pe}_c - \frac{v}{K^{1/2}}\widehat{T}, \tag{S32}$$

where $v := -\mathrm{dPe}_{\mathrm{eff}}/\,\mathrm{d}T|_{T=T_c} > 0$ for an effective Péclet number decreasing through a critical value to trigger the increase in positive feedback. This effective linearisation of $\mathrm{Pe}_{\mathrm{eff}}$ reflects the small change in $\mathrm{Pe}_{\mathrm{eff}}$ across the critical point.

Additionally, we introduce

$$A \sim K^{1/2}\left(B(\widehat{T})\cos\omega z + \frac{A_1}{K^{1/2}}\right), \quad I \sim \frac{\mu r}{\alpha\gamma K^{1/2}}\left(B(\widehat{T})\cos\omega z + \frac{I_1}{K^{1/2}}\right),$$
$$R \sim \frac{r}{\beta}\left(1 + \frac{R_1}{K^{1/2}}\right), \quad C \sim \frac{r}{\gamma K^{1/2}}\left(B(\widehat{T})\cos\omega z + \frac{C_1}{K^{1/2}}\right), \tag{S33}$$

where the asymptotic scalings of $K^{1/2}$ can be justified *a posteriori* and are consistent with the time scaling (S53) and the breakdown of the small $A$ solution (S26).

With the above scalings, the leading-order version of the system (S29) is automatically satisfied, with the scaled amplitude $B(\widehat{T})$ undetermined for now. The remaining goal of this analysis is to determine and analyse the equation that $B$ satisfies, which will inform the dynamic delay in the onset of QS activation. This is achieved by deriving the appropriate governing system for the first-correction terms in the asymptotic expansions of (S33) and deriving an appropriate solvability condition for $B$.

##### C. Solvability condition

At the next asymptotic order, (S29) yields the following system for the first-correction terms

$$\varepsilon K\frac{\mathrm{d}B}{\mathrm{d}\widehat{T}}\cos\omega z = \frac{\partial^2 A_1}{\partial z^2} + \rho\left(q + \frac{r\Lambda}{K}I_1 - rB\cos\omega z\right) - \kappa A_1, \tag{S34a}$$

$$\varepsilon K\frac{\mathrm{d}B}{\mathrm{d}\widehat{T}}\cos\omega z = \alpha\left(C_1 - I_1\right), \tag{S34b}$$

$$0 = -R_1 - B\cos\omega z, \tag{S34c}$$

$$\varepsilon K\frac{\mathrm{d}B}{\mathrm{d}\widehat{T}}\cos\omega z = \left(k_- + \gamma\right)\left(A_1 + R_1 B\cos\omega z - C_1\right), \tag{S34d}$$

with the following boundary conditions from (S30)

$$\frac{\partial A_1}{\partial z} = 0 \quad \text{on } z = 0, \qquad \frac{\partial A_1}{\partial z} + \mathrm{Pe}_c A_1 = v\widehat{T}B\cos\omega \quad \text{on } z = 1. \tag{S35}$$

The system (S34) can be rearranged to obtain the following single ODE for $A_1$:

$$\frac{\partial^2 A_1}{\partial z^2} + \omega^2 A_1 = F\left(z, \widehat{T}; B, \frac{\mathrm{d}B}{\mathrm{d}\widehat{T}}\right), \tag{S36a}$$

where

$$F\left(z, \widehat{T}; B, \frac{\mathrm{d}B}{\mathrm{d}\widehat{T}}\right) := \varepsilon K \cos\omega z \left[1 + \frac{r\rho\Lambda}{K}\left(\frac{1}{\alpha} + \frac{1}{\gamma + k_-}\right)\right]\frac{\mathrm{d}B}{\mathrm{d}\widehat{T}} - \rho q + \rho r B \cos\omega z + \frac{r\rho\Lambda}{K} B^2 \cos^2 \omega z. \tag{S36b}$$

Invoking the Fredholm Alternative Theorem (FAT), the system (S35)–(S36) only admits solutions for particular functions $F$. This is the appropriate solvability condition required to determine $B(\widehat{T})$ in terms of a first-order ODE. We apply FAT in the standard manner: by multiplying the ODE (S36) by $\cos\omega z$, the solution to the homogeneous adjoint problem, and using integration by parts to shift the linear operator onto the adjoint solution. This procedure yields the following solvability condition

$$\int_0^1 F \cos\omega z \,\mathrm{d}z = v\widehat{T} B \cos^2 \omega, \tag{S37}$$

where the right-hand side arises due to the boundary condition (S35) at $z = 1$.

To obtain the appropriate ODE for $B$, we re-write (S37) as

$$\varepsilon K I_2 \left[1 + \frac{r\rho\Lambda}{K}\left(\frac{1}{\alpha} + \frac{1}{\gamma + k_-}\right)\right]\frac{\mathrm{d}B}{\mathrm{d}\widehat{T}} = \rho q I_1 + \left(v\widehat{T}\cos^2\omega - \rho r I_2\right) B - \frac{\rho r \Lambda I_3}{K} B^2, \tag{S38}$$

where we have defined

$$I_n = \int_0^1 \cos^n \omega z \,\mathrm{d}z, \tag{S39a}$$

so that

$$I_1 = \frac{\sin\omega}{\omega}, \quad I_2 = \frac{1}{2} + \frac{\sin 2\omega}{4\omega}, \quad I_3 = \frac{\sin\omega}{\omega} - \frac{\sin^3\omega}{3\omega}. \tag{S39b}$$

The appropriate 'initial condition' for the ODE (S38) arises through asymptotic matching[1] with the quasi-steady solution (S26) as the critical point is approached. This results in the following matching condition

$$B \sim -\frac{\rho q I_1}{v\widehat{T}\cos^2\omega}, \quad \text{as } \widehat{T} \to -\infty. \tag{S40}$$

The ODE (S38) encodes all of the important information about travelling through the bifurcation. In particular, it contains the information about any delay caused by the dynamic effect of crossing the critical point. To transform the ODE into a form more amenable to systematic interrogation, we introduce a rescaled amplitude $\overline{B}(\sigma)$ and time $\sigma$ through the following scalings:

$$B = \sqrt{\frac{qI_1K}{r\Lambda I_3}}\,\overline{B}, \quad \widehat{T} = \frac{\rho r I_2}{v\cos^2\omega} + \frac{\nu_1}{v\sqrt{2\nu_2}}\sigma, \tag{S41}$$

where

$$\nu_1 := \sqrt{\frac{2I_2}{\cos^2\omega}\left[1 + \frac{r\rho\Lambda}{K}\left(\frac{1}{\alpha} + \frac{1}{\gamma + k_-}\right)\right]}, \quad \nu_2 := \frac{I_2 K^2 \cos^2\omega}{\rho^2 q r I_1 I_3 \Lambda}\left[1 + \frac{r\rho\Lambda}{K}\left(\frac{1}{\alpha} + \frac{1}{\gamma + k_-}\right)\right], \tag{S42}$$

defined for later convenience. These scalings transform the ODE (S38) and the initial condition (S59) into

$$\varepsilon v \nu_2 \frac{\mathrm{d}\overline{B}}{\mathrm{d}\sigma} = 1 + \sigma\overline{B} - \overline{B}^2, \qquad \overline{B} \sim -\frac{1}{\sigma}, \quad \text{as } \sigma \to -\infty, \tag{S43}$$

the behaviour of which, importantly, is only controlled by the single parameter grouping $\Omega := \varepsilon v \nu_2$. As such, we can immediately note that if $\Omega \ll 1$ then the system is quasi-steady as we pass through the critical point, and hence there is no discernible lag time in the system. In this case, the form of the right-hand side of the ODE in (S43) reveals that this critical point is an imperfect transcritical bifurcation, where the additional solution branch is negative and therefore unphysical. The imperfect nature of this arises from the base production of autoinducer - if this were zero the bifurcation would be a perfect one. However, in practice $\Omega$ is large and investigating this limit through an asymptotic analysis allows us to understand the delay in the system.

##### D. Quantifying the system lag

When $\Omega := \varepsilon v \nu_2 \gg 1$, the system (S43) evolves over several different timescales, most importantly: $\sigma = O(\sqrt{\Omega})$ where $\overline{B} = O(1/\sqrt{\Omega})$, and $\sigma - \sqrt{2\Omega \log \Omega} = O(\sqrt{\Omega/\log \Omega})$ where $\overline{B} = O(\sqrt{\Omega \log \Omega})$. In the first timescale, setting $\sigma = \sqrt{\Omega} s$ and $\overline{B} = Y/\sqrt{\Omega}$ in (S43), and taking the limit $\Omega \to \infty$, we obtain the leading-order equation

$$\frac{\mathrm{d}Y}{\mathrm{d}s} = 1 + sY, \qquad Y \sim -\frac{1}{s}, \quad \text{as } s \to -\infty, \tag{S44}$$

where we have neglected a small quadratic term in $Y$ of $O(1/\Omega)$. The system (S44) is solved by

$$Y \sim \sqrt{\frac{\pi}{2}} \exp(s^2/2) \left( \operatorname{erf} \frac{s}{\sqrt{2}} + 1 \right), \tag{S45}$$

where erf is the standard error function. The asymptotic solution (S45) is no longer valid once $Y = O(\Omega s)$ for large $s$, at which point the error term is no longer small compared to the terms retained in (S44). This occurs when $s = \sqrt{2 \log \Omega}$. At this point, there is an (interior) boundary layer in time, over which $\overline{B}$ jumps to being of $O(\sqrt{\Omega})$ and after this interior layer the leading-order asymptotic system is governed by

$$\Omega \frac{\mathrm{d}\overline{B}}{\mathrm{d}\sigma} = \sigma \overline{B} - \overline{B}^2, \tag{S46}$$

at leading order, noting that $\sigma = O(\sqrt{\Omega})$. While (S46) can be solved exactly by matching with the interior boundary layer, the important information from this analysis has already been determined. That is, the sudden increase in $\overline{B}$ (and $Y$) occurs when $\sigma = \sqrt{2\Omega \log \Omega}\,(1 + o(1))$. As such, we are able to calculate the delay in the system as we quickly pass through this critical point. That is, for large $\Omega$ the dynamic system will exhibit a jump at time $\sigma = \sqrt{2\Omega \log \Omega}$. Reversing all our scalings into the original slow time $T$, we find that this delay time is

$$T_{\text{delay}} = \frac{\rho r I_2}{v\sqrt{K}\cos^2 \omega} + \nu_1 \sqrt{\frac{\varepsilon \log \varepsilon v \nu_2}{v}}, \quad \text{for } \Omega \gg 1. \tag{S47}$$

Since the first term on the right-hand side is much smaller than the second in practice, we only retain the second term when we re-write the delay time in terms of the original time $t = T/\varepsilon$, to obtain:

$$t_{\text{delay}} = \nu_1 \sqrt{\frac{\log \varepsilon v \nu_2}{\varepsilon v}}, \tag{S48}$$

noting that

$$\varepsilon v := -\left.\frac{\mathrm{d}\mathrm{Pe}_{\text{eff}}}{\mathrm{d}t}\right|_{t=t_c} > 0. \tag{S49}$$

#### V. DELAY TIME FOR A GROWING POPULATION

##### A. Away from the critical point

For a growing population, the analysis is similar to that of decreasing flow. To account for a cell growth, we can consider a density $\rho(\delta t)$ that varies in time over a timescale $\tau = \delta t = O(1)$. Given that the timescale of cell growth is hours compared to the timescale of seconds for vertical diffusion, $\delta \ll 1$. As an instructive example, we note that for exponential cell growth the density would have the form

$$\rho = \rho_0 \exp(\delta t), \tag{S50}$$

with doubling time $(\log 2)/\delta$.

In this system, the governing equations are slightly adapted versions of (S29)–(S30), accounting for a varying density rather than a varying flow. As such, the relevant governing equations are

$$\delta\frac{\partial A}{\partial \tau} = \frac{\partial^2 A}{\partial z^2} + \rho(\tau)\left(q + \lambda I - k_+ AR + k_- C\right) - \kappa A, \tag{S51a}$$

$$\delta\frac{\partial I}{\partial \tau} = \mu C - \alpha I, \tag{S51b}$$

$$\delta\frac{\partial R}{\partial \tau} = r - k_+ AR + k_- C - \beta R, \tag{S51c}$$

$$\delta\frac{\partial C}{\partial \tau} = k_+ AR - k_- C - \gamma C, \tag{S51d}$$

with boundary conditions

$$\frac{\partial A}{\partial z} = 0 \quad \text{on } z = 0, \qquad \frac{\partial A}{\partial z} + \mathrm{Pe}_{\mathrm{eff}} A = 0 \quad \text{on } z = 1. \tag{S52}$$

In the same manner as for varying flow, the form of (S51)–(S52) shows that as $\rho$ changes in $\tau$, the dynamic system is quasi-steady. Therefore, before the onset of positive feedback the asymptotic solution to (S51)–(S52) in the limit $\delta \to 0$ is given by (S24)–(S25).

#### B. Near to the critical point

As with the varying flow problem, this solution breaks down near the onset of positive feedback, which we say occurs at $\tau = \tau_c$, where $\rho(\tau_c) = \rho_c$, as defined implicitly by (S27). Near this breakdown, the dynamics are critically slowed and become more important to the system. To explore these dynamics, we introduce a new timescale

$$\tau = \tau_c + \frac{\widehat{\tau}}{K^{1/2}}, \tag{S53}$$

so the density can be written asymptotically as

$$\rho \sim \rho_c + \frac{u}{K^{1/2}}\widehat{\tau}, \tag{S54}$$

where $u := \mathrm{d}\rho/\,\mathrm{d}\tau|_{\tau=\tau_c} > 0$ for growing cells.

We introduce the asymptotic expansions

$$A \sim K^{1/2}\left(M(\widehat{\tau})\cos\omega z + \frac{A_1}{K^{1/2}}\right), \quad I \sim \frac{\mu r}{\alpha\gamma K^{1/2}}\left(M(\widehat{\tau})\cos\omega z + \frac{I_1}{K^{1/2}}\right),$$
$$R \sim \frac{r}{\beta}\left(1 + \frac{R_1}{K^{1/2}}\right), \quad C \sim \frac{r}{\gamma K^{1/2}}\left(M(\widehat{\tau})\cos\omega z + \frac{C_1}{K^{1/2}}\right), \tag{S55}$$

where the scaled amplitude $M(\widehat{\tau})$ is undetermined for now. Proceeding in the same manner as for the varying flow problem, and substituting (S55) into (S51) we obtain the the following system for the first-correction terms

$$\delta K\frac{\mathrm{d}M}{\mathrm{d}\widehat{\tau}}\cos\omega z = \frac{\partial^2 A_1}{\partial z^2} + \rho_c\left(q + \frac{r\Lambda}{K}I_1 - rM\cos\omega z\right) + \frac{ur\Lambda}{K}\widehat{\tau}M\cos\omega z - \kappa A_1, \tag{S56a}$$

$$\delta K\frac{\mathrm{d}M}{\mathrm{d}\widehat{\tau}}\cos\omega z = \alpha\left(C_1 - I_1\right), \tag{S56b}$$

$$0 = -R_1 - M\cos\omega z, \tag{S56c}$$

$$\delta K\frac{\mathrm{d}M}{\mathrm{d}\widehat{\tau}}\cos\omega z = \left(k_- + \gamma\right)\left(A_1 + R_1 B\cos\omega z - C_1\right), \tag{S56d}$$

with boundary conditions

$$\frac{\partial A_1}{\partial z} = 0 \quad \text{on } z = 0, \qquad \frac{\partial A_1}{\partial z} + \mathrm{Pe}_{\mathrm{eff}} A_1 = 0 \quad \text{on } z = 1. \tag{S57}$$

The system (S56)–(S57) yields the solvability condition

$$\delta K I_2 \left[1 + \frac{r\rho_c \Lambda}{K}\left(\frac{1}{\alpha} + \frac{1}{\gamma + k_-}\right)\right] \frac{\mathrm{d}M}{\mathrm{d}\widehat{\tau}} = \rho_c q I_1 + r I_2 \left(\frac{u\Lambda}{K}\widehat{\tau} - \rho_c\right) M - \frac{\rho_c r \Lambda I_3}{K} M^2, \tag{S58}$$

where $I_n$ are defined in (S39).

The appropriate 'initial condition' for the ODE (S58) arises through asymptotic matching[1] with the quasi-steady solution (S26) as the critical point is approached. This results in the following matching condition

$$M \sim -\frac{\rho_c q I_1 K}{r I_2 u \Lambda \widehat{\tau}}, \quad \text{as } \widehat{\tau} \to -\infty. \tag{S59}$$

Importantly, we note that the scalings

$$M = \sqrt{\frac{q I_1 K}{r \Lambda I_3}}\overline{B}, \quad \widehat{\tau} = \frac{\rho_c K}{u\Lambda} + \frac{\rho_c}{u I_2}\sqrt{\frac{q I_1 I_3 K}{r\Lambda}}\sigma, \tag{S60}$$

transform the ODE (S58) into

$$\delta u \nu_3 \frac{\mathrm{d}\overline{B}}{\mathrm{d}\sigma} = 1 + \sigma \overline{B} - \overline{B}^2, \tag{S61}$$

where

$$\nu_3 := \frac{I_2^2 K}{\rho_c^2 q I_1 I_3}\left[1 + \frac{r\rho_c\Lambda}{K}\left(\frac{1}{\alpha} + \frac{1}{\gamma + k_-}\right)\right], \tag{S62}$$

and

$$\delta u := \left.\frac{\mathrm{d}\rho}{\mathrm{d}t}\right|_{t=t_c} > 0. \tag{S63}$$

The ODE (S61) is equivalent to the corresponding rescaled ODE for varying flow, given in (S43), except for a different definition of the constant premultiplying the derivative on the left-hand side. As such, the resulting analysis is equivalent, with an appropriate substitution of constant. Using parameters from Table S1, we show that the typical delay time due to growth is of the same timescale as growth in Fig. S9.

#### VI. SIMULATION PARAMETER SETS

We determined an approximate order of magnitude for each dimensional parameter by taking measured or estimated values in LuxIR-type systems from the literature (see Table S1). In each simulation, all parameters in Table S1 were fixed except cell density $\rho$ and Pe, which were varied in time-dependent simulations as indicated in the main text. In reality, the values of the kinetic parameters will vary widely owing to error in measurements or estimates, and to differences between species and synthetic systems. Therefore, to illustrate that the general predictions in the main text are valid for a wide range of possible parameter values, we also performed the following simulations for Fig. S4:

1. Production terms $\tilde{r}$ and $\tilde{q}$ were all reduced or increased by an order of magnitude, while fixing all other parameters to be their values in Table S1 (with $\tilde{H} = 5$ $\mu$m, $\dot{\gamma} = 1000$ /s, 5000 /s and 10000 /s, and $L = 10$ $\mu$m).

2. Decay terms $\tilde{\alpha}$, $\tilde{\beta}$, $\tilde{\gamma}$ and $\tilde{\kappa}$ were all reduced or increased by an order of magnitude, while fixing all other parameters to be their values in Table S1 (with $\tilde{H} = 5$ $\mu$m, $\dot{\gamma} = 1000$ /s, 5000 /s and 10000 /s, and $L = 10$ $\mu$m).

3. Binding terms $\tilde{k}_-$ and $\tilde{k}_+$ were all reduced or increased by an order of magnitude, while fixing all other parameters to be their values in Table S1 (with $\tilde{H} = 5$ $\mu$m, $\dot{\gamma} = 1000$ /s, 5000 /s and 10000 /s, and $L = 10$ $\mu$m).

4. Feedback terms $\tilde{\lambda}$ and $\tilde{\mu}$ were all reduced or increased by an order of magnitude, while fixing all other parameters to be their values in Table S1 (with $\tilde{H} = 5$ $\mu$m, $\dot{\gamma} = 1000$ /s, 5000 /s and 10000 /s, and $L = 10$ $\mu$m).

5. Diffusion term inside the cell population $\tilde{D}_c$ was reduced by one or two orders of magnitude, while fixing all other parameters to be their values in Table S1 (with $\tilde{H} = 5$ $\mu$m, $\dot{\gamma} = 1000$ /s, 5000 /s and 10000 /s, and $L = 10$ $\mu$m).

| Symbol | Definition | Order of magnitude and units | Reference |
|---|---|---|---|
| $\tilde{H}$ | Height of cell population | Variable (2-10 $\mu$m) | Typical height of early biofilm or cell colony[2] |
| $\dot{\gamma}$ | Shear rate applied to cell population | Variable (100-10,000 /s) | Typical range of shear rates in natural and industrial environments[3] |
| $\tilde{L}$ | Length of cell population in dowstream direction | Variable (4-20 $\mu$m) | Chosen to depend on $\tilde{H}$ such that the height to length aspect ratio is 0.5, as is typical in early biofilms or colonies[2] |
| $\tilde{D}_e$ | Diffusion coefficient of AI in water | 500 $\mu m^2$/s | Approximate typical estimate of AI diffusion in water used in previous studies[4–6] |
| $\tilde{D}_c$ | Diffusion coefficient of AI in cell layer | 500 $\mu m^2$/s | Approximate typical estimate of AI diffusion in water used in previous studies[4–6]. For simplicity, we also use this value in the cell population, although it may be lower depending on the conditions[5] and on AI-matrix interactions (see Fig. S5) |
| $\tilde{A}_0$ | Autoinducer concentration at activation threshold | 5 nM | Taken from various measurements[7–9]. Note that the variable $\tilde{A}_0$ is a free parameter that does not affect the model results; we use this value for convenience, so that A=1 at activation. |
| $\tilde{q}$ | Basal production of AI | 1 nM/s | Assuming production of approximately 1 molecules/s/cell, as estimated in various previous studies[10–12]; we convert to nM using a cell volume of 0.4 $\mu m^3$ (as estimated in Ref. 2) |
| $\tilde{r}$ | Production of LuxR | 0.01 nM/s | Assuming production of approximately 0.01 molecules/s/cell, estimated from transcription and degradation rates of the relevant mRNA, and translation rate, in Ref. 13; we convert to nM as before. |
| $\tilde{\mu}$ | Coefficient of activation of LuxI by C | 1000 /s | By our linearization $\tilde{\mu} \approx \tilde{\mu}_a/\tilde{K}_a$, where $\tilde{\mu}_a$ is the activated production rate (in units of nM/s) and $\tilde{K}_a$ is the dissociation constant or activation coefficient (in units of nM). Using an activated expression rate of LuxI of 1000-10000 nM/s (approximately corresponding to the estimate in Ref. 14; we have converted to nM as before) and a dissociation constant of 1-10 nM (a measurement of $\tilde{K}_a = 10$ nM is given for TraR in Refs. 15,16), we obtain $\tilde{\mu} = 1000$ /s (see Fig. S2 for a discussion of the effect of $K_a$) |
| $\tilde{\lambda}$ | Coefficient of catalysis of AI by LuxI | 0.01/s | Turnover number measured to be approximately 1 /min by Ref. 17 |
| $\tilde{k}_-$ | Unbinding of AI and LuxR | 0.01 /s | Approximately consistent with previous estimates in the literature of 1 /min (see Refs. 10,13,18) |
| $\tilde{k}_+$ | Forward binding of AI and LuxR | 0.0001 /nM/s | Set to give a dissociation constant of 100 nM (as measured in Ref. 19) |
| $\tilde{\beta}$ | Decay of LuxR | 0.01 /s | Taken from measurements of TraR half life of approximately 1 min (see Ref. 16) |
| $\tilde{\gamma}$ | Decay of C | 0.001 /s | Set to be approximately ten times slower than LuxR degradation[16] |
| $\tilde{\alpha}$ | Decay of LuxI | 0.001 /s | Taken to be same order of magnitude as C degradation |
| $\tilde{\kappa}$ | Decay of AI | 0.0001 /s | Approximately consistent with estimated internal and external degradation rates of 0.01/min (thought to be dependent on pH)[10,20–22] |
| $\rho$ | Volume fraction of cells in population | Function of time (typical value 0.1) | Measured number density of cells at onset of quorum sensing was approximately 0.25 cells / $\mu m^3$ in Ref. 23. Using typical cell volume of 0.4 $\mu m^3$ (measured by Ref. 2) gives volume density of 0.1 |

TABLE S1: Estimated typical orders of magnitude of the dimensional parameters in the problem. Where possible, we take orders of magnitude consistent with measured values in the literature. Otherwise, we use values in which previous studies fitted simulations to data, or we use typical estimates from the literature. These parameters could vary widely between species or in synthetic applications and should not be taken to be definitive values, but are used to guide our understanding of the relevance of the various identified asymptotic regimes.

The parameter sets used in all simulations are available via Github at https://github.com/philip-pearce/quorum-flow.

---


[1] van Dyke, M. *Perturbation methods in fluid dynamics* (Parabolic Press, 1975).

[2] Hartmann, R. *et al.* Emergence of three-dimensional order and structure in growing biofilms. *Nature Physics* **15**, 251–256 (2019).

[3] Marcos, Fu, H. C., Powers, T. R. & Stocker, R. Bacterial rheotaxis. *Proceedings of the National Academy of Sciences* **109**, 4780–4785 (2012).

[4] Stewart, P. S. Diffusion in Biofilms. *Journal of Bacteriology* **185**, 1485–1491 (2003).

[5] Horswill, A. R., Stoodley, P., Stewart, P. S. & Parsek, M. R. The effect of the chemical, biological, and physical environment on quorum sensing in structured microbial communities. *Analytical and Bioanalytical Chemistry* **387**, 371–380 (2007).

[6] Emge, P. *et al.* Resilience of bacterial quorum sensing against fluid flow. *Scientific Reports* **6**, 33115 (2016).

[7] Kaplan, H. B. & Greenberg, E. P. Diffusion of autoinducer is involved in regulation of the Vibrio fischeri luminescence system. *Journal of Bacteriology* **163**, 1210–1214 (1985).

[8] Andersen, J. B. *et al.* gfp-Based N-Acyl Homoserine-Lactone Sensor Systems for Detection of Bacterial Communication. *Applied and Environmental Microbiology* **67**, 575–585 (2001).

[9] Collins, C. H., Arnold, F. H. & Leadbetter, J. R. Directed evolution of Vibrio fischeri LuxR for increased sensitivity to a broad spectrum of acyl-homoserine lactones. *Molecular Microbiology* **55**, 712–723 (2005).

[10] Tanouchi, Y., Tu, D., Kim, J. & You, L. Noise reduction by diffusional dissipation in a minimal quorum sensing motif. *PLoS Computational Biology* **4**, 4–11 (2008).

[11] Alberghini, S. *et al.* Consequences of relative cellular positioning on quorum sensing and bacterial cell-to-cell communication. *FEMS Microbiology Letters* **292**, 149–161 (2009).

[12] Leaman, E. J., Geuther, B. Q. & Behkam, B. Quantitative Investigation of the Role of Intra-/Intercellular Dynamics in Bacterial Quorum Sensing. *ACS Synthetic Biology* **7**, 1030–1042 (2018).

[13] Goryachev, A., Toh, D. & Lee, T. Systems analysis of a quorum sensing network: Design constraints imposed by the functional requirements, network topology and kinetic constants. *Biosystems* **83**, 178–187 (2006).

[14] Müller, J., Kuttler, C., Hense, B. A., Rothballer, M. & Hartmann, A. Cell–cell communication by quorum sensing and dimension-reduction. *Journal of Mathematical Biology* **53**, 672–702 (2006).

[15] Zhu, J. & Winans, S. C. Autoinducer binding by the quorum-sensing regulator TraR increases affinity for target promoters in vitro and decreases TraR turnover rates in whole cells. *Proceedings of the National Academy of Sciences* **96**, 4832–4837 (1999).

[16] Zhu, J. & Winans, S. C. The quorum-sensing transcriptional regulator TraR requires its cognate signaling ligand for protein folding, protease resistance, and dimerization. *Proceedings of the National Academy of Sciences* **98**, 1507–1512 (2001).

[17] Schaefer, A. L., Val, D. L., Hanzelka, B. L., Cronan, J. E. & Greenberg, E. P. Generation of cell-to-cell signals in quorum sensing: Acyl homoserine lactone synthase activity of a purified Vibrio fischeri LuxI protein. *Proceedings of the National Academy of Sciences of the United States of America* **93**, 9505–9509 (1996).

[18] Weber, M. & Buceta, J. Dynamics of the quorum sensing switch: Stochastic and non-stationary effects. *BMC Systems Biology* **7**, 6 (2013).

[19] Urbanowski, M. L., Lostroh, C. P. & Greenberg, E. P. Reversible Acyl-Homoserine Lactone Binding to Purified Vibrio fischeri LuxR Protein. *Microbiology* **186**, 631–637 (2004).

[20] Schaefer, A. L., Hanzelka, B. L., Parsek, M. R. & Greenberg, E. P. Detection, purification, and structural elucidation of the acylhomoserine lactone inducer of Vibrio fischeri luminescence and other related molecules. *Methods in Enzymology* **305**, 288–301 (2000).

[21] Basu, S., Gerchman, Y., Collins, C. H., Arnold, F. H. & Weiss, R. A synthetic multicellular system for programmed pattern formation. *Nature* **434**, 1130–1134 (2005).

[22] Kaufmann, G. F. *et al.* Revisiting quorum sensing: Discovery of additional chemical and biological functions for 3-oxo-N-acylhomoserine lactones. *Proceedings of the National Academy of Sciences of the United States of America* **102**, 309–314 (2005).

[23] Singh, P. K. *et al.* Vibrio cholerae Combines Individual and Collective Sensing to Trigger Biofilm Dispersal. *Current Biology* **27**, 3359–3366.e7 (2017).

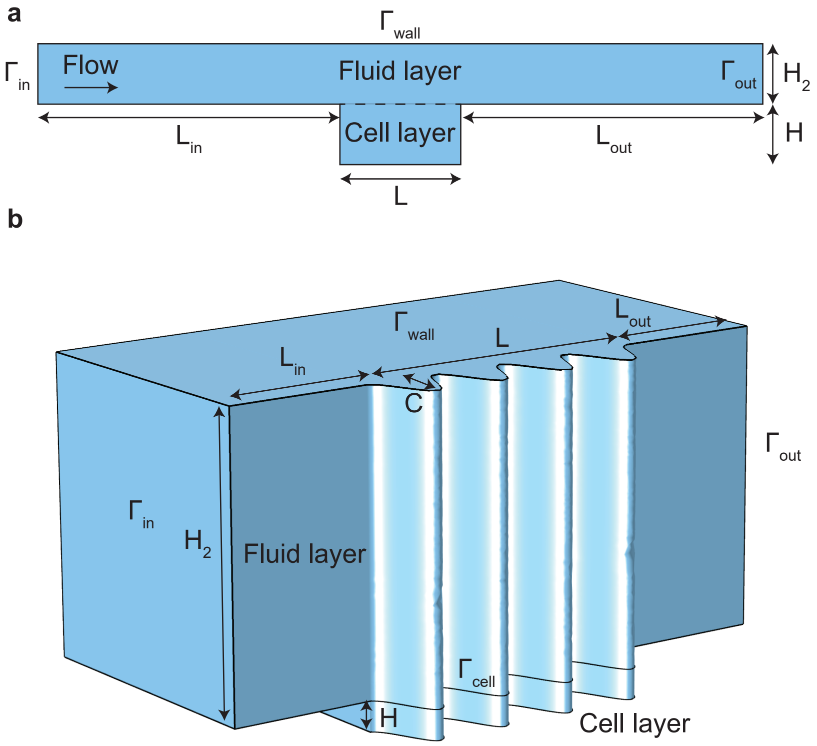


FIG. S1: Schematic diagrams of simulation domains and boundaries. Our numerical procedure was as follows. The problem is solved numerically in Comsol Multiphysics 5.5 on two domains: the flow layer and the cell layer. We include an inflow and outflow region in the flow layer to ensure the boundary conditions do not impose artificial physics upon the results. a) In 2D simulations, the velocity field in the fluid layer is imposed as a uniform constant-shear flow. Eq. (4) (see main text) is solved in the fluid layer for autoinducer concentration $A$. In the cell layer, where it is assumed that no flow is present, Eqs. (1)-(2) (see main text) are solved for the concentration of the autoinducers and LuxIR proteins. As boundary conditions on autoinducer concentration $A$, zero concentration, $A = 0$, is imposed on the inflow boundary, $\Gamma_{\text{in}}$, and zero flux, $\mathbf{n} \cdot \nabla A = 0$, is imposed on all other external boundaries, $\Gamma_{\text{wall}}$ and $\Gamma_{\text{out}}$. On the internal boundary between the cell layer and the fluid layer (dashed line) continuity of concentration $A$ and its derivatives is imposed. In unsteady simulations, the initial condition is taken to be the steady solution for the mean Péclet number, which also corresponds to the value of the Péclet number at time $t = 0$. The lengths of the inflow and outflow regions, $L_{\text{in}}$ and $L_{\text{out}}$, and the height of the fluid layer, $H_2$, are taken to be large enough such that the boundary conditions do not affect the results. For $L_{\text{in}}$ and $L_{\text{out}}$, a length 5 times the height of the cell layer was found to be sufficient. For $H_2$, it was found to be sufficient to ensure that $LD/\dot{\gamma}H_2^2 < 0.1$. b) In 3D simulations, the fluid layer has height $H_2 = 50\ \mu$m and the cell layer that coats the floor has height $H = 5\ \mu$m. The crevices are described by the curve $d = C\left((-1 + \cos(4\pi s/10))^2\right)$ where $C$ is the dimensionless crevice depth and $s$ is the dimensionless distance in the downstream direction (both scaled by $H$), and $s = 0$ is located at the upstream edge of the cell layer. The velocity field $\mathbf{u}$ in the fluid layer is obtained by solving Eq. (4) (see main text), with the viscosity and density taken to be those of water. Zero pressure, $p = 0$, is imposed on $\Gamma_{\text{out}}$, and the pressure on $\Gamma_{\text{in}}$, $p = p_{\text{in}}$, is imposed such that the maximum shear rate applied to the cell layer is approximately 1000 $\text{s}^{-1}$ (thus the channel Reynolds number is approximately 0.25). A no-slip condition, $\mathbf{u} = 0$, is applied at the remaining external boundaries $\Gamma_{\text{wall}}$, and at the boundary between the cell layer and the fluid layer, $\Gamma_{\text{cell}}$. The concentrations of autoinducers and LuxIR proteins are then solved for as before. All code required to generate the simulation results is available on Github at https://github.com/philip-pearce/quorum-flow (requires Comsol 5.5 license, Matlab license, and Comsol LiveLink with Matlab License).

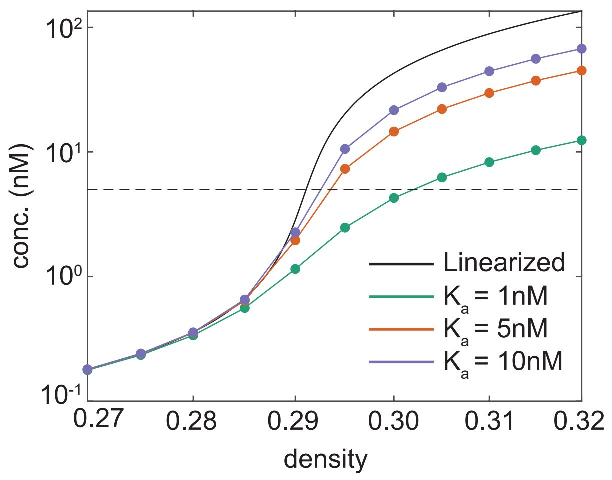


FIG. S2: Effect of reduction in LuxI activation via saturation in autoinducer-LuxR dimer binding to DNA on quorum sensing (QS) activation across the critical density. The maximum concentration of autoinducers in the population is plotted for 2D simulations with the typical parameters listed in Table S1 (and $\tilde{H} = 5\ \mu$m, $\dot{\gamma} = 1000\ \text{s}^{-1}$, $L = 10\ \mu$m). The black line shows the results for a linearized activation term $\tilde{\mu}\tilde{C}$ in Eq. (S2), and the remaining lines show the results for a nonlinear activation term $\tilde{\mu}_a\tilde{C}/\left(\tilde{K}_a + \tilde{C}\right)$ in Eq. (S2). In the nonlinear simulations, we vary the dissociation constant $\tilde{K}_a$ of the dimers from the DNA; for clarity, we fix the ratio $\tilde{\mu}_a/\tilde{K}_a$, i.e. the linearized activation parameter $\tilde{\mu}$ (see Table S1). A measure of the nonlinearity or effect of saturation can be obtained by comparing the concentration of dimers $\tilde{C}_{\text{act}} \approx r\tilde{A}/\gamma K$ (by the steady solution Eq. S21, since $K$ is typically large) at the onset of activation, $\tilde{A} = 5$ nM, to the dissociation constant $\tilde{K}_a$. If $\tilde{C}_{\text{act}}/\tilde{K}_a$ is large, we expect saturation in binding to DNA at the onset of activation; in this case, the nonlinearities in the activation term will be important, and the concentration of autoinducers at activation will be reduced from the linearized case. If $\tilde{C}_{\text{act}}/\tilde{K}_a$ is small, we do not expect saturation in binding to DNA at the onset of activation; in this case, the concentration of autoinducers at activation will be close to the linearized case. Using the dissociation constant of 10 nM measured in Ref. 15, we find $\tilde{C}_{\text{act}}/\tilde{K}_a \approx 0.05$ nM/10nM $= 0.005$, which explains why the linearized approximation is successful at capturing the onset of QS activation for these parameters (purple line). For systems with smaller dissociation constants, the estimate of the critical density is still quite accurate, but the maximum concentration after QS activation is lowered; therefore, in such systems the population-level robustness of QS activation would be expected to be reduced.

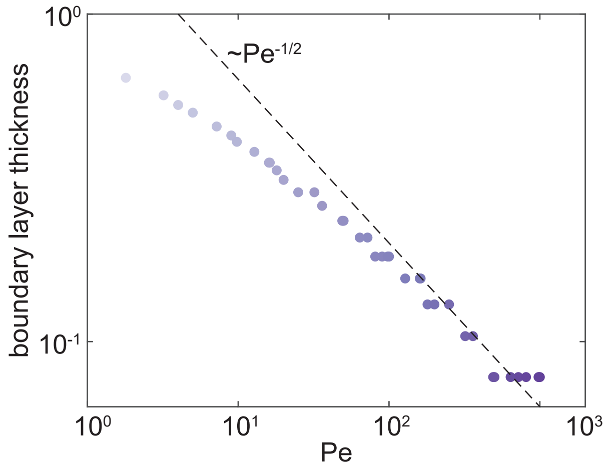


FIG. S3: The thin diffusive boundary layer at the downstream edge of the cell population has a non-dimensional thickness (scaled by the population height) which is $O(\text{Pe}^{-1/2})$. The Péclet number is plotted against the distance from the downstream edge of the location where the calculated effective Péclet number at the boundary, $A_y/A$, is minimised. Simulations correspond to those in Fig. 2 in the main text (for clarity, colors correspond to the value of the Péclet number in the same way as in Fig. 2). The dashed line corresponds to an arbitrary constant multiplied by $\text{Pe}^{-1/2}$, for comparison.

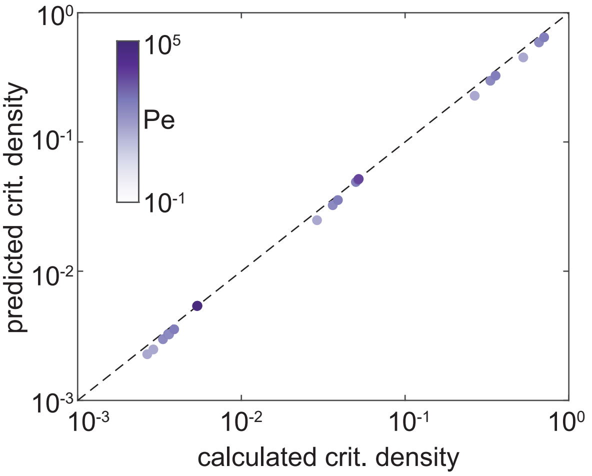


FIG. S4: Calculated versus predicted critical densities for a range of kinetic and physical parameters. Simulations were performed as described in Fig. 1 in the main text, but with parameters given in the 'Simulation Parameter Sets' section of the Supporting Information. Note that some points are indistinguishable because at high Péclet number the critical density does not depend strongly on the shear rate.

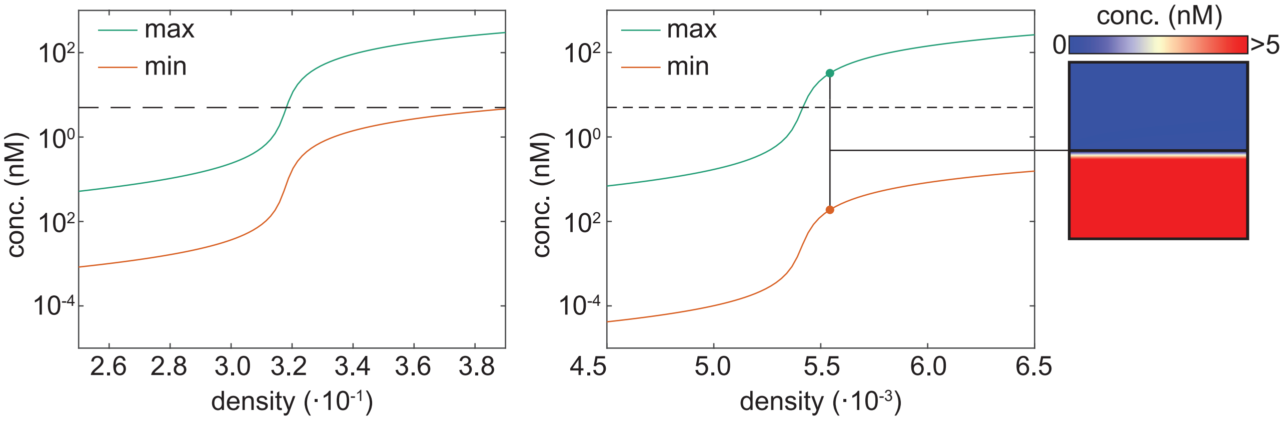


FIG. S5: Effect on steady QS activation of a reduced diffusion coefficient in the cell population (so that $D > 1$; see Eq. S9), which can be caused by interactions between autoinducers and the extracellular matrix. Simulations correspond to those in Fig. 1c in the main paper, with $\dot{\gamma} = 10000\ \mathrm{s}^{-1}$, and with the diffusion coefficient inside the cell population reduced by a) a factor of 10 ($\tilde{D}_c = 50\ \mathrm{m}^2/\mathrm{s}$) and b) a factor of 100 ($\tilde{D}_c = 5\ \mathrm{m}^2/\mathrm{s}$). We set a) $H = 2\ \mu\mathrm{m}$ and b) $H = 5\ \mu\mathrm{m}$; the latter plot is for comparison with the bottom panel of Fig. 1c in the main paper. Compared to corresponding populations but with identical diffusion coefficients, a reduction in diffusion inside the population causes an increase in the effective Péclet number and a reduction in the critical density through Eqs. (6)-(8) in the main paper (compare panel b to the bottom panel of Fig. 1c). Furthermore, owing to the increased effective Péclet number, a decreased $D$ increases spatial heterogeneity inside the cell population compared to the situation where $D$ is equal in the cell population and the flow, although QS activation remains robust - even for the very high effective Péclet numbers shown in panel b, only a very small region of the cell population near the boundary between the population and the flow remains inactivated for densities above the critical value (inset).

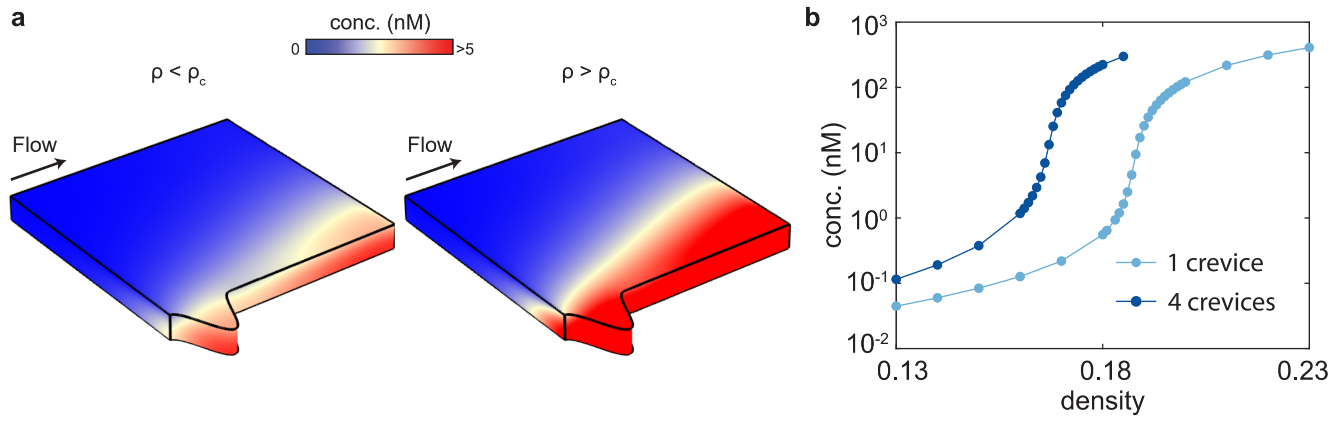


FIG. S6: The effect of the number of crevices on steady quorum sensing activation (simulations correspond to those in Fig. 3, but in a channel with a single crevice instead of four). a) Autoinducer concentration fields for a density just below the activation threshold (left, $\rho = 0.187$) and just above the activation threshold (right, $\rho = 0.188$) show that quorum sensing is first activated in the crevice and in the downstream region of the population. After activation, the whole downstream region becomes activated. b) A cell population in a channel with a single crevice requires a higher cell density for activation than a population in a similar geometry but with multiple crevices (corresponding to the geometry shown in Fig. 3).

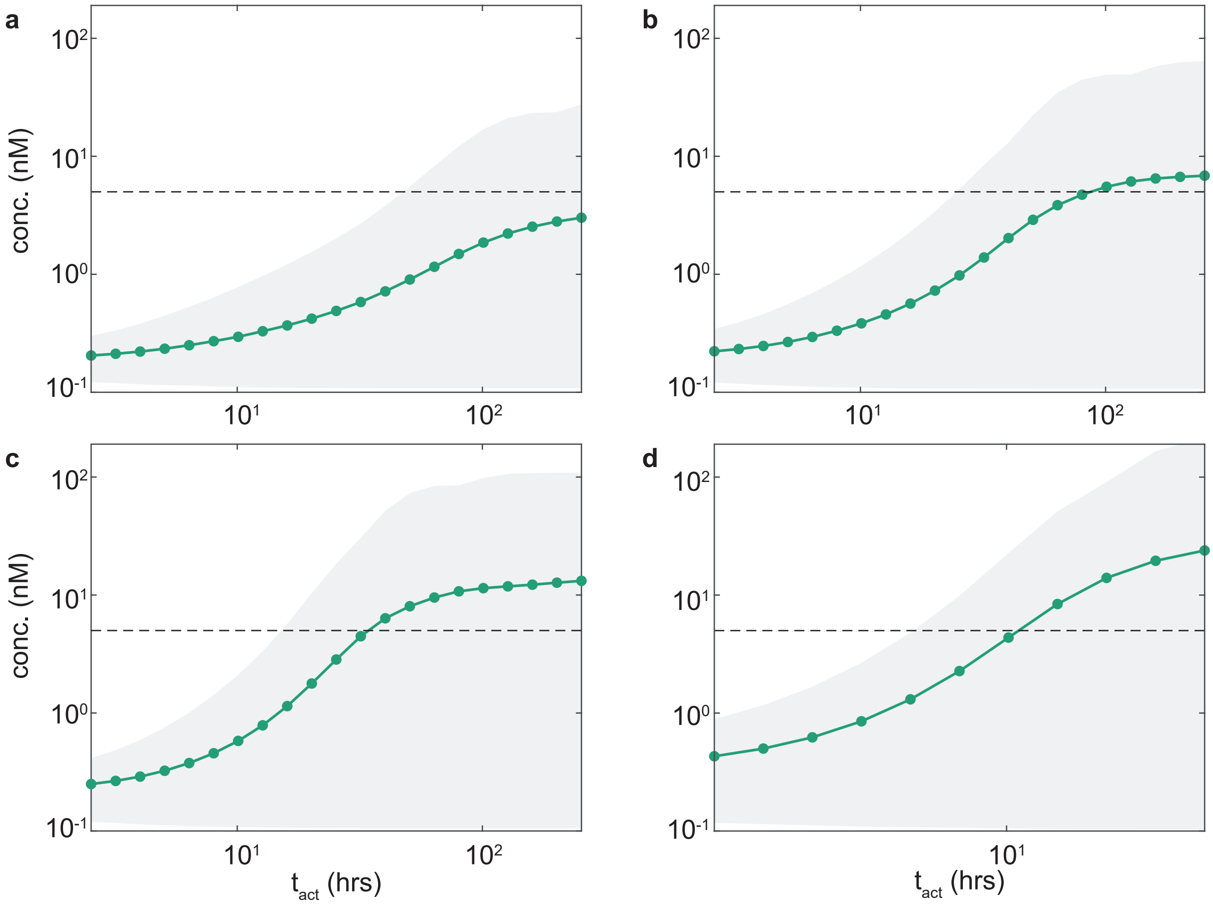


FIG. S7: The critical activation time for an oscillating flow, defined as the value of $t_{\mathrm{act}}$ for which the range of the maximum concentration (grey area) first passes above the activation threshold $\tilde{A} = 5$ nM, is found to be of order of magnitude 10 hours across a range of oscillations, as predicted in the main text. Simulations correspond to the same kinetic parameters and mean shear rate as those shown in Fig. 4b in the main text, but with an amplitude in the oscillations of a) 7500 $\mathrm{s}^{-1}$, b) 8000 $\mathrm{s}^{-1}$, c) 8500 $\mathrm{s}^{-1}$, d) 9500 $\mathrm{s}^{-1}$. Dynamic QS activation is found to be weaker for oscillations that enter into the activation region of parameter space for a smaller proportion of the total oscillation period (panels a and b).

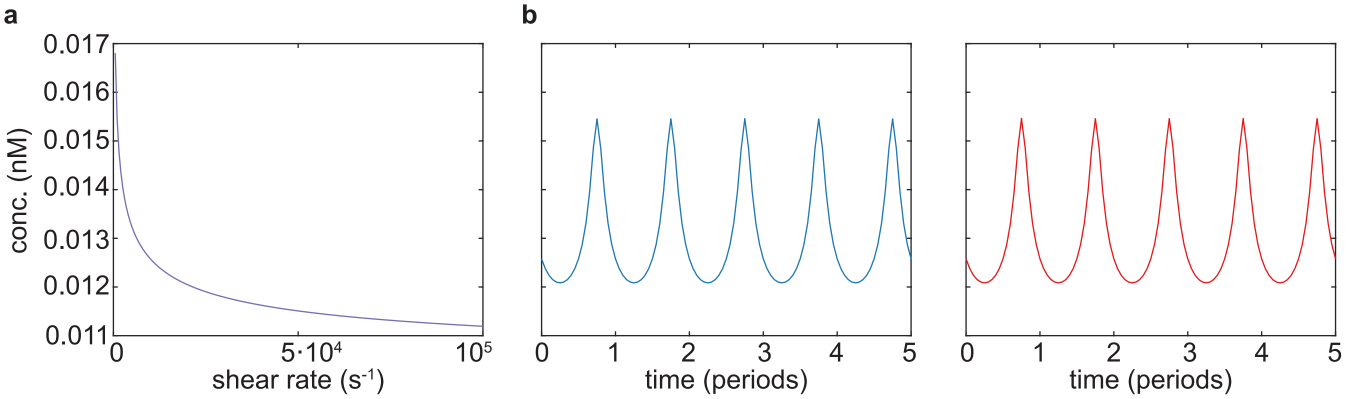


FIG. S8: Shear rate oscillations in a system without feedback (simulations correspond to those in Fig. 4, but with $\lambda = 0$; we note that the binding parameters $k_+$ and $k_-$ do not have a distinguishable effect in this case). a) In steady flow, the autoinducer concentration does not vary over orders of magnitude for the range of shear rates studied (corresponding to the ones shown in Fig. 4). b) In unsteady flow, the system behaves almost identically for slow oscillations (left, $t_{\text{act}} =$, corresponding to the blue line in Fig. 4b inset) and fast oscillations (right, $t_{\text{act}} =$, corresponding to the red line in Fig. 4b inset). The range of concentration values on the vertical axis is the same in each panel.

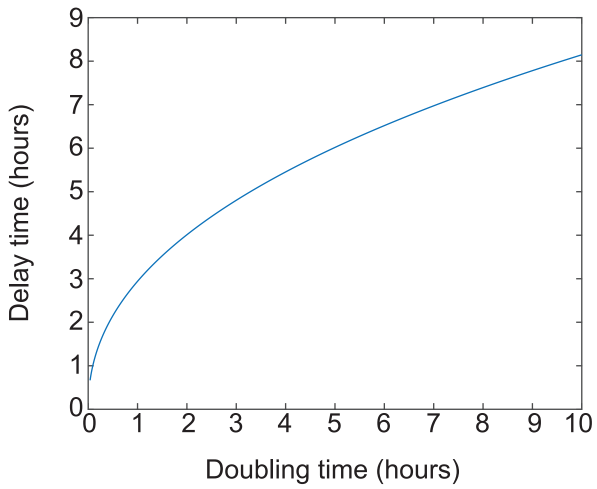


FIG. S9: The delay time associated with population growth is of the same order of magnitude as the doubling time, for the kinetic parameters listed in Table S1. The delay time was calculated using the large-$\Omega$ results of Section IV D, but with $\Omega = \delta u \nu_3$, as described in (S62)–(S63).

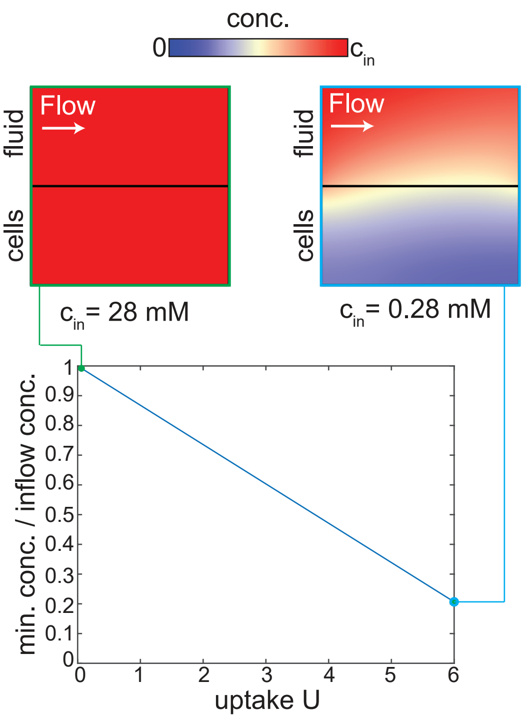


FIG. S10: We expect nutrients to be plentiful in the conditions considered in this study, so that all cells can be assumed to be physiologically active. This is based on the observation that cells in biofilms with radius around 10 $\mu$m remain uniformly growth-active even in shear rates a hundredfold smaller than the smallest shear rates used here[2] (where a smaller shear rate implies reduced access to nutrients). To confirm this assumption further, we perform simulations of a nutrient that satisfies an advection-diffusion equation in the flow, and a diffusion-reaction equation in the cell layer, where the reaction rate is taken to be an uptake rate multiplied by the cell density. For simplicity, we use parameters for glucose estimated in Ref. 23. Therefore, we take the diffusion coefficient to be $D = 600\ \mu\mathrm{m}^2/\mathrm{s}$ in both the fluid layer and the cell layer, and the uptake rate in the cell layer to be constant at $10^8$ molecules/min/cell, or approximately $u = 10^7$ nM/s (we convert to nM/s using the same assumptions as in Table S1). We set the cell layer height to be $H = 10\ \mu$m (the largest population size used in this study), and the channel height to be $H_2 = 100\ \mu$m, using the same geometry as in Fig. S1a. We take the cell density to be 0.1 (see Table S1), and the shear rate to be $\dot{\gamma} = 100\ \mathrm{s}^{-1}$, which corresponds to the smallest shear rate used in this study, and therefore the most nutrient-limiting conditions. The system is characterised by the Péclet number $\mathrm{Pe} = \dot{\gamma}H^2/D$ and a non-dimensional uptake rate $U = uH^2/Dc_{\mathrm{in}}$, where $c_{\mathrm{in}}$ is the nutrient inflow concentration. We find nutrients to be plentiful for a range of values of $U$ from $U = 0.06$ up to $U = 6$, which spans values of $c_{\mathrm{in}}$ from $c_{\mathrm{in}} = 28$ mM (corresponding to the glucose concentration of M9 medium with 0.5% glucose[23]) to $c_{\mathrm{in}} = 0.28$ mM. For populations with significantly reduced access to nutrients, the assumption of uniform base autoinducer production throughout the population may need to be relaxed.
